# Predicting social experience from dyadic interaction dynamics: the BallGame, a novel paradigm to study social engagement

**DOI:** 10.1101/2024.01.26.577035

**Authors:** Annika Lübbert, Malte Sengelmann, Katrin Heimann, Till R. Schneider, Andreas K. Engel, Florian Göschl

## Abstract

Theories of embodied cognition suggest that a shared environment and ongoing sensorimotor interaction are central for interpersonal learning and engagement. To investigate the embodied, distributed and hence dynamically unfolding nature of social cognitive capacities, we present a novel laboratory-based coordination task: the BallGame. Our paradigm requires continuous sensing and acting between two players who jointly steer a virtual ball around obstacles towards as many targets as possible. By analysing highly resolved measures of movement coordination and gaming behaviour, game-concurrent experience ratings, semi-structured interviews, and personality questionnaires, we reveal contributions from different levels of observation on social experience. In particular, successful coordination (number of targets collected) and intermittent periods of high versus low movement coordination (variability of relation) emerged as prominent predictors of social experience. Importantly, having the same (but incomplete) view on the game environment strengthened interpersonal coordination, whereas complementary views enhanced engagement and tended to generate more complex interactive behaviour. Overall, we find evidence for a critical balance between similarity and synchrony on the one hand, and variability and difference on the other, for successful engagement in social interactions. Finally, following participant reports, we highlight how interpersonal experience emerges from specific histories of coordination that are closely related to the interaction context in both space and time.

## Introduction: social cognition involves interactions that span across levels of organisation

When humans collaborate to solve a problem, a myriad of things happens. The environment shapes the language, movements and social roles we have available and choose from. The specifics of the task bring certain routines and skills to the foreground. Our personality, as well as our self-confidence and general condition influence our expectations, experience and behaviour.

The complex set of processes at work during collaborative action has inspired a diverse audience of researchers. Here, we present an experimental design and analysis approach that serves the integration of several perspectives on social interaction research. More specifically, we present a task that engages two participants in an interactive computer game, and perform analyses that integrate their gaming behaviour, finger movement coordination, subjective experience and personality traits. At the heart of our approach is the interest in relationality: how do two players co-determine their interaction dynamics? How do different elements of this process, such as personality differences, the interaction context, players’ performance levels or their degree of movement coordination, relate?

Our approach is directly inspired by recent proposals to ground social cognition in interactive sensorimotor coordination (Varela et al., 1991; Clark, 1997; Menary, 2010; Engel et al., 2013). The concept of ‘socialising sensorimotor contingencies’ in particular (Lübbert et al., 2021) highlights sensing and acting in mutual response as the key organising principle of social cognition. In this regard, we take a pragmatic stance: we locate social cognition in the domain of relationships between individuals and describe social behaviour and experience as the consequence of dynamic cycles of informational and sensorimotor coupling between agents. To bring this perspective into the cognitive science laboratory, we test here whether changes in the experienced quality of interaction are associated with changes in sensorimotor coordination between interacting players. As reviewed by Lübbert and colleagues (2021), empirical studies from dance and music to classical cognitive science laboratory settings have linked movement synchronisation to neural synchronisation of interacting individuals (Dumas et al., 2010; Zhou et al., 2016), to their subjective experience (Llobera et al., 2016; Jakubowski et al., 2020; Ramseyer & Tschacher, 2016), as well as to contextual factors such as individual differences or task constraints (Feniger-Schaal et al., 2018; Vesper et al., 2016). However, studies that make room for interactive autonomy, generate detailed records across more than two levels of observation, consider changes in time and interaction context, and bridge domains by integrating approaches and findings, remain scarce. To contribute to their development, we present an experimental setting that combines an engaging interactive task with multiple forms of qualitative and quantitative observation: the BallGame.

In the BallGame, two players jointly steer a virtual ball around obstacles and towards as many targets as possible. The BallGame offers participants possibilities for action that are overlapping (both players can steer the ball in any direction with equal maximal force), diverse (at any moment there are many possible ways forward) and stimulating (the game control and task are neither too easy nor too difficult, and present collaborative advantages). Because we are interested in sensorimotor contingencies as a substrate of social cognition, we chose to include continuous movement and ongoing gaming dynamics (instead of discrete actions such as button presses and coordination through turn taking). Additionally, we used a game controller that is unfamiliar to most people: it required steering a virtual ball by bending and stretching one’s index fingers. This allowed participants to start at the same level of experience. Finally, we included trials in our experiment that featured obstacles visible only to one of the two players. This condition both challenged and stimulated interpersonal coordination because players accessed different but overall more information.

Besides recording participants’ finger movements and gaming behaviour, we asked them to rate their experience in terms of their perceived level of ball control, engagement, agreement with and predictability of their partner. Participants also self-assessed their personality traits using questionnaires, and we performed individual interviews at the end of each experiment.

By investigating individual experience as an interactive property - a characteristic of ongoing sensorimotor, interpersonal and situated action - our design reflects current trends towards relationality in the cognitive sciences (O’Regan & Noë, 2001; De Jaegher & Di Paolo, 2007; Konvalinka & Roepstorff, 2012; Clark, 2016; Durt et al., 2017; Lübbert et al., 2021). These strands of research urge us to locate social cognition at interrelating and intersecting levels of organisation: from biological to cultural factors, in individuals, interacting parties as well as their environment. This implies that empirical investigation of social cognitive processes should consider dynamics across multiple levels of observation. We believe that our approach meets this demand. Our participants needed to master a challenging game control (precise index finger movements) and had to coordinate their steering actions with a partner, both of which stimulates engagement and creates room for individual choice. We further considered changes in behaviour and social experience over different periods of time, and assessed the influence of seeing the same versus in part different obstacles compared to one’s partner. In our principal line of investigation, we then predicted participants’ social experience from a combination of multiple operationalisations of interpersonal movement coordination, gaming behaviour, personality differences as well as the interaction context. In line with the concept of socialising sensorimotor contingencies, we hence investigated social cognition as a process that establishes and details itself in embodied and situated action.

The central research question that we pursued with the present study focuses on the relationship between social experience and interpersonal sensorimotor coordination: is social experience (partly) constituted by how we move with our interaction partner? Can we, thus, use measures of interpersonal movement coordination to predict how participants experience their interaction? Our second line of investigation concerns the evolution of participants’ interaction over time and across conditions of joint play. In particular, we tracked changes over blocks (3-4 minutes of play) and sessions (20 minutes), and tested for differences in social behaviour and experience at times when participants had the same or partially different views on the game environment. Finally, prompted by unexpected findings in the interviews, we investigated individual differences at the transition from joint back to individual play, as well as the within*-*trial evolution of the interaction dynamics.

## Methods

### Participants

23 pairs of players (14 female-female pairs, 8 male-male pairs, 1 female-male pair; mean age 24.7 years, range 20-37) participated in the BallGame. Participants received monetary compensation for their time and a bonus depending on their success at the game (0.7 cents per collected target). Participants had normal or corrected to normal vision and reported no history of neurological or psychiatric illness. The study was approved by the Ethics Committee of the Medical Association Hamburg and conducted in accordance with the Declaration of Helsinki. Prior to the recordings, all participants provided written informed consent.

### The BallGame

We designed the BallGame in order to facilitate social engagement, continuous interaction and participant autonomy, while ensuring rigorous multi-level game-concurrent observation. The BallGame is a computer-based task in which two players use their index fingers to steer a virtual ball across a two-dimensional surface, avoiding obstacle regions to collect as many targets as possible in limited time (*Figure 1A, left* presents a screenshot of the game environment). Each trial of the game lasted one minute. At any point in a trial, three targets and six obstacles were visible to each player, and three additional obstacles remained invisible. Players could learn about the location of invisible obstacles by keeping track of areas in which the ball reliably slowed down. When players collected a target (when the ball hit one of the three visible coins), this target disappeared, and the previously inactive (fourth) target appeared. The challenge was to learn to steer the ball (first alone, then together), get to know the landscape (remember the location of invisible obstacles) and collect as many targets as possible in limited time: after each one-minute trial, the constellation of obstacles and targets shifted. The three outer targets (visible in *Figure 1A, left*) rotated around the equidistant centre, and another 9 of 15 possible obstacle locations were activated (six visible, three invisible). The 9 obstacle locations were pseudo-randomly picked from 15 possible locations, so that all direct lines between the targets were blocked by at least one obstacle. There were 60 different surfaces for all pairs (with the order of the landscapes shuffled within subsequent blocks of the same game condition).

**Figure 1.**
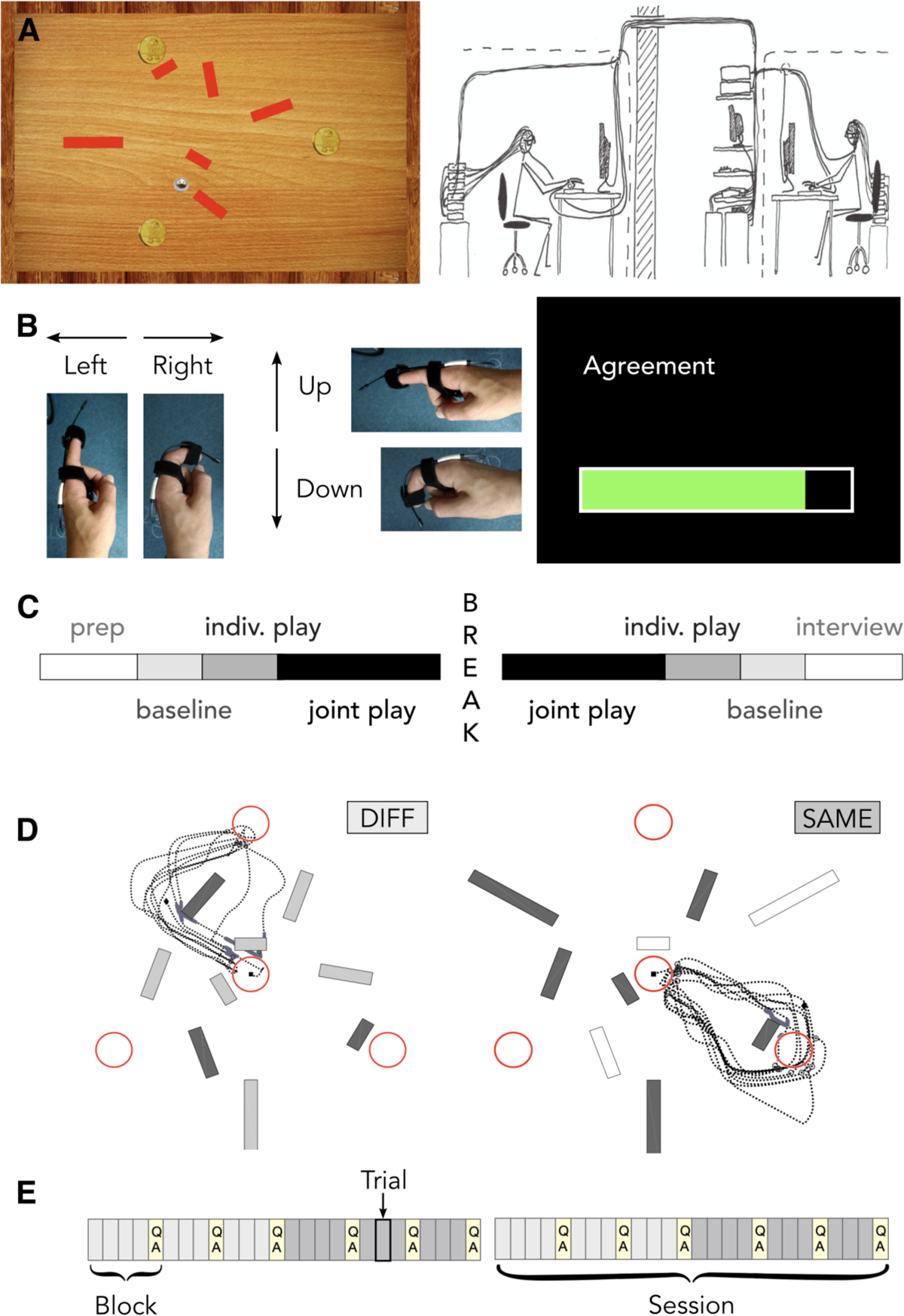
Experimental Paradigm. **(A, left)** Screenshot of the game environment. Participants steered the ball (grey marble) to collect targets (golden coins) and avoid obstacles (red bars that slow down the ball to 10% of its speed). **(A, right)** Illustration of the BallGame setup: a pair of participants, each equipped with a 128-channel EEG cap, eye-tracker goggles and bimetal sensors attached to index fingers, sitting in adjacent rooms / EEG chambers. **(B, left)** Demonstration of the game control - a bimetal sensor attached to the index finger translated bending and stretching of the finger into ball movement on the screen. **(B, right)** View of an example experience rating (bar filled by ‘left-right’ movement, answer confirmed with long ‘down’ movement). **(C)** Experimental protocol. After the instructions, participants were prepared (prep) for the game-concurrent data recording. The experiment began with baseline tasks and 10 trials of individual play. After further 20 trials of joint play, we took a longer break. Afterwards, participants played another 20 trials of joint, and 10 trials of individual play, and completed the baseline tasks. The experimentended with individual interviews, during which the other participant filled in personality questionnaires. **(D)** The two joint play conditions. In joint play DIFF ( different), three of n ine obstacles were visible to both players (dark grey bars), three only to player one *or* two (light grey bars). In joint play SAME, players saw the same six of nine obstacles (dark grey bars) - three obstacles remained invisible to the team (empty bars). The black dotted line indicates the path traveled by the ball in an example one-minute trial. **(E)** Experimental protocol of the joint play period. Joint play was structured in 12 blocks of three or four trials each, after which participants rated their experience in terms of their level of engagement, agreement and predictability (light-yellow boxes marked ‘QA’ (questions and answers)). In each session, participants play ed 10 trials of each condition (light grey boxes = joint play DIFF; dark grey boxes = joint play SAME).

Throughout a trial, participants continuously influenced the movement of the ball, with either index finger controlling the acceleration of the ball along the x and y axis, respectively. During joint play, players’ acceleration was accumulated (up to a maximal speed), such that the ball quickly moved right when both players steered right, slowly to the right when players steered at orthogonal directions centred around rightward movement, and not at all, when players’ steering directions were opposite. Though prompted by our interest in (continuous) social sensorimotor contingencies, this design feature was particularly inspired by research with a highly reduced space for dyadic interaction: the perceptual crossing paradigm (Auvray & Rohde, 2012; Froese et al., 2014). In this setting, two players move an avatar across a digital line and receive a stimulus (e.g., a vibration on their finger tip) each time they encounter the other, the other’s shadow or a stationary object. This scenario leads players into stable sensorimotor interaction dynamics, allowing them to reliably detect each other’s presence. Findings from the perceptual crossing paradigm convinced us that players could identify their partner’s actions (within overlapping, continuous game control) and learn to coordinate.

Over the course of the experiment, participants played the BallGame in three different conditions: individual play, joint play with the same obstacle visibility (SAME) and joint play with in part different obstacle visibility (DIFF; see *Figure 1D* for an illustration of the two joint play conditions). Beyond the parallel with natural social engagement (in which interacting partners hold complementary views and information), this design feature was inspired by Vesper and colleagues’ (2016) findings about the strong influence of shared perceptual information on how individuals accomplish coordinated action. Overall, joint play presented a collaborative advantage in the form of cumulative acceleration (though maximal ball speed is the same in joint and individual play) and complementary information (during joint play DIFF). However, players also needed to differentiate hitting an invisible obstacle from disagreeing with their partner (steering in opposite directions), which was particularly challenging during joint play DIFF, where unilaterally (in)visible obstacles were presented.

### Experimental protocol

Participants were scheduled to arrive at the institute at the same time. When both participants had finished reading the written game instructions, the experimenter orally summarised the most important points and provided further information about the procedure and game environment, including a reminder of the collaborative advantage: in half of the joint-play trials (joint play ‘DIFF’ condition, see *Figure 1D*), their partner would see the three obstacles that remained invisible to themselves. Since participants knew neither of the experimental structure (*Figure 1E*) nor which joint play condition they were currently playing, it was advisable for them to always coordinate with their partner, that is to pay attention to their steering directions as potential signals for invisible obstacles.

After clarifying remaining questions, participants took their seats in the two EEG chambers, situated in adjacent rooms (see *Figure 1A, right*). With a team of one to three assistants, the experimenter then prepared the game-concurrent data collection: participants were equipped with 128-channel passive electrode EEG caps (EASY CAP BC-128-x7, Herrsching, Germany) to record their brain activity, eye-tracker goggles (Pupil Core, Pupil Labs, Germany) to trace their pupil dilation and gaze-fixation, and bimetal sensors (Finger Twitch Transducer SS61L, BIOPAC Systems, USA) at both index fingers, used as game-control and to answer the questions about their experience of the game (see *Figure 1B*). The eye-tracker and bimetal sensors were then calibrated to fit individual movement ranges. After these preparations, participants completed baseline tasks intended to serve as localisers for later EEG analyses (*note that the present work does not include analyses of the EEG and eye tracking data*). Next, participants performed 10 trials of individual play to familiarise themselves with the BallGame, in particular the game control. We then proceeded with four times 10 trials of joint play, with the order of conditions (joint play SAME and DIFF) balanced over pairs. Afterwards, participants played alone again and completed another round of the baseline tasks. Halfway through the joint play period, we took a longer break during which participants could relax, use the bathroom, stretch or step outside. See *Figure 1C* for an overview of the experimental protocol. During the play period, we informed participants about transitions between individual and joint play, and asked them to rate their experience every three to five trials (see below, *Levels of Observation*). We invited participants to use these moments as small breaks. *Figure 1E* illustrates the experimental protocol of the joint play period. After completing the experiment, we conducted a semi-structured interview with each participant about their experiences playing the game. While one participant was interviewed, the other filled in personality questionnaires.

### Levels of observation

To capture the ongoing interaction dynamics during the BallGame, we organised our analysis along four *levels of observation*: personality traits, experience, gaming behaviour and finger movement. Below, we describe how we measured and parametrised activity at each level - *Figure 2* gives an overview of all parameters considered in the present analysis.

*Temporal resolution:* For all measures except personality traits, we assessed changes over time: across sessions (first vs. second half of joint play), blocks (the first four, second three and last three trials played under the same obstacle visibility condition) and, wherever possible, trial segments (3 x 20 seconds). There was a short break between the sessions, implying that participants actually experienced a first and a second part of the game. Blocks ran in parallel with the intervals at which participants rated their experience - we hence aggregated data from the three or four trials that preceded a rating. Finally, the rationale for splitting each trial into three segments derived from our findings in the interviews (see below, *Results - Within-trial changes in the gaming dynamic*).

**Figure 2.**
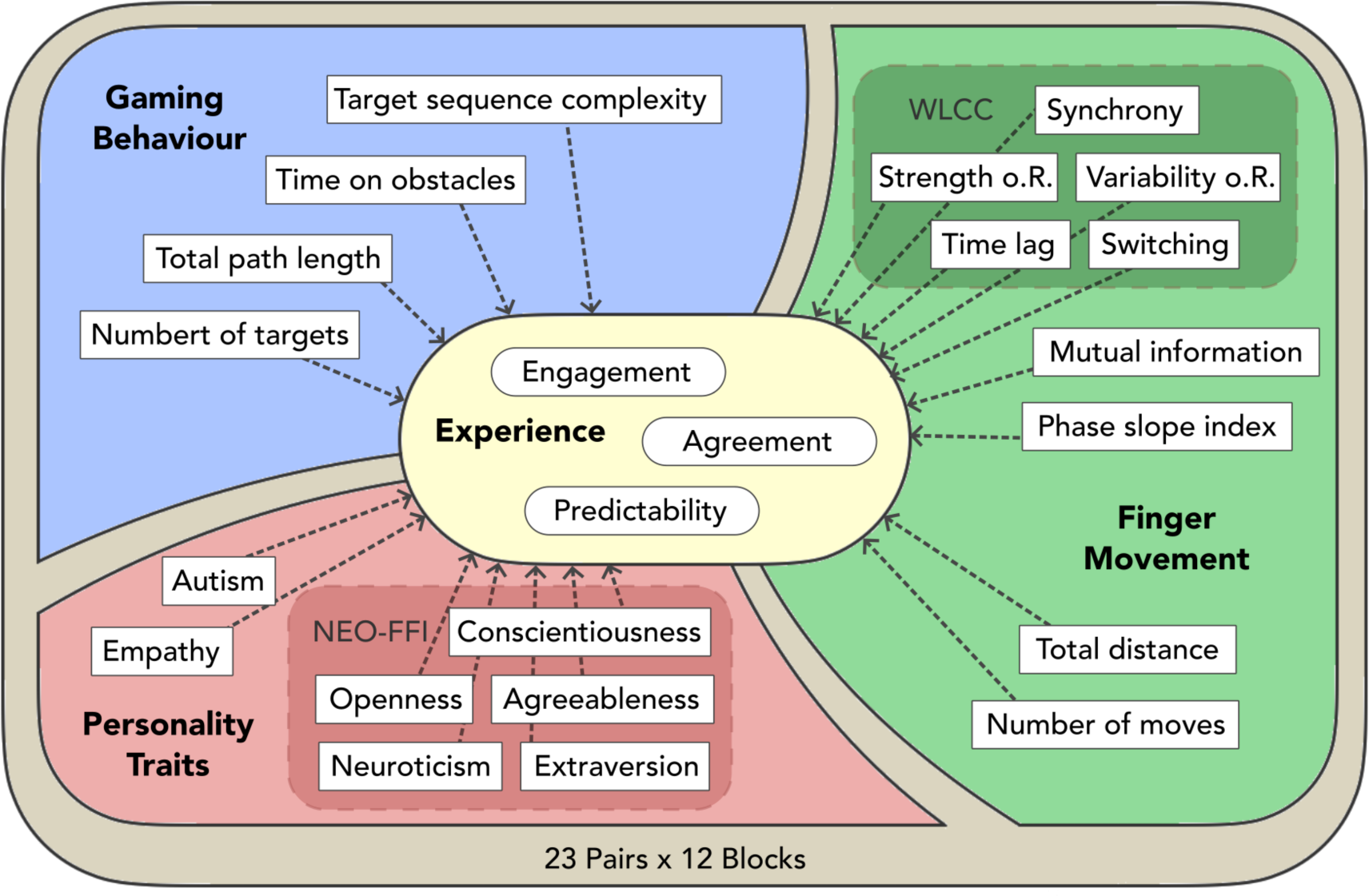
Predicting experience from multiple levels of observation: parameters assessed in the BallGame. We considered four levels of observation of interpersonal coordination: personality traits (**red**), gaming behaviour (**blue**), finger movement (**green**) and experience (**light-yellow**). Each level is described through several parameters. Light-shaded boxes indicate a family relationship between parameters: within personality, this concerns the five traits assessed by the NEO five factor inventory (NEO-FFI); within finger-movement, this concerns five measures of coordination derived from a windowed lagged cross-correlation analysis (WLCC).

*Personality traits*: participants filled in the NEO-FFI (Borkenau & Ostendorf, 2008), a general personality questionnaire that allows self-description along the dimensions of neuroticism, extraversion, openness to experience, agreeableness, and conscientiousness. They further completed the Autism Quotient (Baron-Cohen et al., 2001), and the SPF-IRI (Paulus, 2009), an interpersonal reactivity index that differentiates four subcomponents (perspective-taking, fantasy, empathic concern, and personal distress) which we aggregated (excluding the last factor) as our ‘Empathy’ measure.

*Experience ratings*: at fixed moments during the game - that is after trials 5, 10 (individual play); 14, 17, 20; 24, 27, 30; 34, 37, 40; 44, 47, 50 (joint play); 55 and 60 (individual play) - participants provided experience ratings. These ratings assessed whether participants felt able to steer the ball through their finger movements (**ball control** - only during individual play), how focused and involved they were in the game (**engagement** - throughout the entire play period), their sense of agreement and smooth performance with their partner (**agreement** - only during joint play), as well as whether they felt they understood what their partner was doing (**predictability** - only during joint play). Participants used the game control (the bimetal sensors attached to their index fingers) to provide their answers through a continuous slider. We translated their rating into integers from 0 to 100. After assessing the distributions of the rating data, we used the raw experience ratings for ball control, agreement and predictability ratings, but transformed the engagement ratings using the Arcsine transformation.

*Participant interviews:* We conducted a semi-structured individual interview with each participant at the end of the experiment. This allowed us to systematically and thoroughly assess the nature of participant engagement in the BallGame. We then opened the interview with generic questions (“What comes to mind when you think back to playing the BallGame?”, “Which moments, if any, were exhausting/fun/social?”), in order to avoid biasing participants. After that, we turned to specific aspects of the game, asking questions that directly relate to our research interests (“Was your partner present to you? If so, when and how?”, “On a scale from 0 = ‘100% PC game’ to 10 = ‘100% social interaction’, how did you experience joint play?”). The full interview sheet is provided in *Supplementary Materials B*.

*Thematic content of the interviews:* We performed a thematic content analysis of the individual post-game interviews, following Elo and Kyngäs (2008) and Kuckartz (2012). Accordingly, we inductively developed a coding scheme that was tested by means of an iterative coding and refining procedure until a quarter of the data could be classified completely and unambiguously. We then continued to code the remainder of the dataset, occasionally merging or refining codes to avoid very small categories (containing less than 5 of the 46 individuals), or to accommodate novel content.

*Gaming behaviour*: we used four parameters to capture participants’ gaming behaviour (see also *Figure 2*) - generating one value per pair during the joint play, and separate values for each player during the individual play period. (1) **Number of targets collected**: for each third of a trial (i.e. 20 seconds), we counted the number of targets collected. (2) **Time spent on obstacle regions**: for each third of a trial, we divided the number of frames the ball spent on any of the obstacles by the total number of frames. (3) **Total path length**: for each third of a trial, we calculated the total distance covered by the ball. (4) **Target sequence complexity**: for each trial, we evaluated how many times the ball went back and forth between two targets. That is, we counted target collection events that did not involve going back and forth between the same two targets, and divided by the total number of targets collected in this trial. The resulting ‘complexity index’ ranges between 0 and 1, with lower values indicating a tendency to stick to a once identified path.

*Finger movement (basics)*: we calculated two basic movement properties. (1) **Movement**: to generate a simple measure that captures the overall amount of finger movement, we integrated the velocity of both fingers, regardless of the direction of movement, for each trial segment. (2) **Number of moves** (direction changes): to estimate the stability of steering, we counted how many times participants switched direction on the x- or y-axis in each third of a trial.

*Finger movement (coordination)*: to quantify the degree of coordination between participants’ finger movements, we calculated seven parameters that assess either the relation between players’ movements (undirected coordination), or potential leader-follower dynamics (directed coordination). All parameters are calculated based on participants’ combined x- and y-axis movement, that is, the angle into which players steered the ball (‘steering direction’).

Our first set of measures is based on a *windowed lagged cross-correlation* (WLCC) analysis, in which we calculated the Spearman correlation between participants’ steering direction over short windows of time. In line with Moulder and colleagues (2018), we generated five measures of coordination: we quantified (1) **synchrony** as the average WLCC coefficient across all lags (see *Supplementary Materials A.1* for WLCC parameters), (2) **strength of relation** as the mean peak-picked WLCC (ppWLCC) coefficient (the largest coefficient of correlation closest to a lag of zero), independent of the lag at which it was observed, (3) **variability of relation** as the standard deviation across ppWLCC coefficients, (4) **time lag** as the average absolute ppWLCC lag (ignoring which participant led or lagged, showing only the relative time delay between players’ steering directions). Finally, we assessed (5) **switching** behaviour as the standard deviation over ppWLCC lags. To control for similarities in movement that may have been induced by the game landscape, we calculated surrogate levels of synchrony: here, we used data of players from different pairs that were navigating the same game landscape (see *Figure S.1*).

We further quantified mutual information (**MI**) and calculated the phase slope index (**PSI**; Nolte et al., 2008) between players’ steering directions. MI quantifies the mutual dependence between two signals and denotes the reduction of uncertainty about one signal that can be achieved by observing the other (Cohen, 2014; Quian Quiroga & Panzeri, 2009). PSI is a measure that quantifies the direction of information flow in multivariate time series. Formally, it corresponds to the weighted average of the slope of the phase of cross-spectra between two signals. In our case, these two signals are the steering directions of two players jointly steering a ball.

See *Supplementary Materials A* for a more detailed introduction of our measures of movement coordination.

### Statistical Analyses

*Predicting social experience (linear mixed effects models)*: to test whether participants’ experience ratings can be predicted from finger movement coordination, gaming behaviour and inter-personal differences, we calculated three linear mixed effects models (using R packages ‘lme4’, Bates et al., 2015, and ‘lmerTest’, Kuznetsova, Brockhoff & Christensen, 2017), one for each of our three social experience ratings (engagement, agreement and predictability - always taking the mean value of both players’ answers). In parallel with participants’ experience ratings, we aggregated all data into 12 blocks, yielding 276 observations per measure (23 pairs x 12 blocks). We initiated each model with the complete set of predictors (4 measures of gaming behaviour, 9 measures of finger movement and 7 measures of personality difference as fixed main effects, no interactions were included), a random intercept for pairs, a time parameter that continuously models the 12 blocks of the joint play period, and an autoregressive covariance structure that models the temporal dependence of repeated measures by allowing for greater similarity of observations that are closer in time (de Haan-Rietdijk, Kuppens & Hamaker, 2016). *Figure 2* illustrates the initial model. We then used a restricted maximum-likelihood estimator to fit the model and iteratively eliminated non-significant predictors until only significant predictors were left. This hierarchical backwards elimination procedure was not applied to the random intercept and the autoregressive covariance structure. Furthermore, we performed a leave-one-out cross-validation procedure to test the generalisability of our findings: we calculated a repeated measures correlation (using R package ‘rmcorr’, Bakdash & Marusich, 2021; see also, Bakdash & Marusich, 2017) between the actual (mean) ratings of our players, and the ratings we predicted based on model parameters that were fit to data from all but the present pair. Following the same rationale, we also calculated pair average correlations. In both cases, higher correlations between observed and predicted ratings indicate better generalisability of the model. Note, however, that this procedure only considered fixed effects.

*Variance over time and across game conditions (MANOVAs and ANOVAs)*: to test for general trends in our game-concurrent observations (experience, gaming behaviour, and finger movement), we calculated multivariate repeated measures analyses of variances (MANOVAs) (using the R package ‘MANOVA.RM’, Friedrich, Konietschke & Pauly, 2021) with three within-pair factors for each family: session (before vs. after the break in the middle of joint play), condition (SAME vs. DIFFerent obstacle visibility) and block (accumulating data in parallel with the intervals at which we ask questions). We determined p-values based on parametric bootstrapping and calculated the modified ANOVA-type statistics (MATS) that can account for potential heteroscedasticity as well as singular covariance matrices, thus relaxing the assumptions of the model, and providing more reliable results with small sample sizes (Friedrich & Pauly, 2018). Below, we report MATS instead of parametric statistics such as the F value. Note that in the ANOVA, we tested for differences between the three subsequent blocks played under the same game condition. In our mixed effects models of participants’ experience ratings, in turn, our time parameter considered changes across all 12 subsequent blocks of joint play.

#### Follow-up analyses in response to unexpected findings from the interviews

Conducting the interviews extended our understanding of how participants played the BallGame. Based on the thematic content analysis, we learned that participants’ experience of the last period of individual play diverged drastically - while some felt relieved of the burden of having to coordinate with their partner, others lost the motivation to play. There were furthermore specific moments in which their interaction partner tended to be especially present to participants: right before and at the beginning of a trial. Finally, the objects in the game environment were omnipresent in participants’ reports about their social experience. To learn more about the interaction dynamic as highlighted by participants, we then conducted three follow-up analyses of our game-concurrent measures of observation:

*Individual differences after the transition from joint to individual play*: To investigate the differences in participant reports about the second period of individual play, we looked for within-group differences in our measures of observation and used the degree of coordination to split our group of participants into two. We classified pairs as strongly versus weakly coordinated based on their aggregate rank on all seven measures of movement coordination (excluding the median pair from this analysis). We then compared the behaviour and experience of strongly versus weekly coordinated players as they shift from joint back to individual play. First, we performed a repeated measures ANOVA of the number of targets collected during the second session with mode of play (individual vs. joint) as within-pairs, and coordination level as between-pairs factors. As before, we determined p-values based on parametric bootstrapping and calculated the ANOVA-type statistics (ATS). In addition to performance, we looked at experience ratings of the final 10 trials of individual play: do sense of ball control or engagement evolve differently in the two coordination groups? Here, the ANOVA compared sense of control ratings in the early versus late individual play period (ball control was not assessed during joint play), and engagement ratings during joint versus individual play of the second session.

*Within-trial changes in the gaming dynamics*: To follow up on participants’ reports of differently experiencing the early versus later parts of a trial, we calculated a repeated measures MANOVA with trial segment as the only within-pairs factor for both movement coordination and gaming behavioural measures, followed by individual measure ANOVAs. Note that several measures of observation were excluded from this analysis, because of insufficient or unavailable data at the level of the trial third, namely: experience ratings, target sequence complexity, PSI and MI.

*Coordination as a function of target and obstacle proximity*: Prompted by participants frequent mention of objects in the game environment, we related the strength of relation (see above) to the time that has passed since the last target was collected - that is, over the target collection cycle. For each moment of the ppWLCC calculation, we identified the fraction of frames that have passed until the next target is collected. We then calculated the mean and standard error of the strength of relation between participants’ finger movements in 20 sub-sections with the same number of entries along the target collection cycle - beginning and ending at the moment a target is collected. We did so separately for the two joint play conditions (SAME versus DIFF), and tested for difference between the conditions in each of the 20 segments along the target cycle, correcting for multiple comparisons using the Benjamini-Hochberg approach (Benjamini & Hochberg, 1995; Haynes, 2013) to control false discovery rates (FDR) across the number of bins. We further assessed the influence of nearby obstacles on coordination: for each moment of the ppWLCC, we determined the visibility of the obstacle that was closest to the ball (minimal distance to the borders of any of the nine obstacles active on the current trial), that is, whether the obstacle was visible to both, either or none of the players. We then performed a repeated measures ANOVA of strength of relation with obstacle-visibility and game condition as within-pairs factors.

*Post-hoc tests & correction for multiple comparisons (ANOVAs).* When we observed a significant effect of the factors block or trial segment (both of which are three-stepped), we used the MANOVA.RM R-package to calculate post-hoc comparisons between individual blocks or trial segments. To correct for multiple comparisons, we used the Benjamini-Hochberg approach (Benjamini & Hochberg, 1995; Haynes, 2013) to control false discovery rates (FDR). Note that we corrected in five groups: one ‘meta group’ formed by the four MANOVAs that aggregate measures within each level of observation (A. experience ratings, B. gaming behaviour, C. basic finger movement, and D. finger movement coordination), and four ‘sub-groups’, each accounting for all ANOVAs and post hoc paired comparisons that we calculated *within* a given level of observation (number of effects = number of parameters at a given level of observation * 7 effects [3 main effects + 3 two-way interactions + 1 three-way interaction] + possible post hoc comparisons for significant effects of block or trial segment]).

## Results

Our analyses integrated multiple levels of observation of continuous, engaged social interaction dynamics and focused in particular on interpersonal sensorimotor coordination as a predictor of social experience. This approach was motivated by recent proposals to ground social cognition in interpersonal coupling mechanisms (Lübbert et al., 2021). Overall, our results demonstrated that social experience in the BallGame was influenced by variables from each of our levels of observation: gaming behaviour (especially the number of targets), movement coordination (in particular the variability of relation between players), personality differences and the interaction context (joint play SAME vs. DIFF, time, objects in the game environment).

To illustrate the nature of social interaction in the BallGame, we begin with an overview of participant reports. We then present our findings from the linear mixed effects models and analyses of variance grouped along three major themes: (1) predictors of social experience; (2) learning effects: changes in gaming behaviour and finger movement parameters over blocks and sessions; and (3) differences between the joint play conditions: effects of seeing the same versus different obstacles.

Finally, prompted by unexpected findings from the interviews, we present results from follow-up analyses. These focus on: (A) individual differences at the transition from joint to individual play, (B) within-trial changes in the gaming dynamics, and (C) game objects as attractors of attention.

### Participant reports: social interaction in the BallGame

The thematic content analysis of participant reports revealed seven major themes (see *Figure 3*): game environment, positive emotion, negative emotion, social presence, strategy, individual play and technical comments, each made up of several sub-codes. *Figure 3* illustrates how many participants talked about a given code. For a complete summary of the interview contents, consult *Supplementary Materials B.2*.

Importantly, the interviews revealed a strong social focus: participants were concerned with figuring out what the partner sees or intends to do (ibid, theme ‘*strategy*’, sub-category ‘listening where to go’, n = 25 participants), in particular during the early trial period (sub-category of ‘listening where to go’, n = 10 participants). Relatedly, participants frequently reported reflecting on whether it was disagreement with their partner or encounter with an invisible obstacle that caused the ball to slow down (theme ‘*social presence*’, sub-category ‘us or obstacle’, n = 19 participants). They also described moments in which difficulties were resolved as particularly pleasant and social (theme ‘*positive emotion*’, sub-categories ‘challenge’ & ‘joint play’, n = 9 participants). What is more, a group of participants experienced a need to re-learn the game control when switching from joint to individual play, possibly indicating strong interpersonal attunement (theme ‘*individual play*’, sub-category ‘readjustment’, n = 14 participants). When asked explicitly, participants also rated their experience of the BallGame as a social interaction rather than a computer game (0 = PC game, 10 = social interaction; mean = 6.45, standard deviation = 1.35). Finally, the interviews confirmed the above mentioned learning effects: participants reportedly learned to coordinate better over time, both concerning steering the ball, as well as interacting with their partner (theme ‘*technical comments*’, sub-category ‘over time’, n = 14 participants).

**Figure 3.**
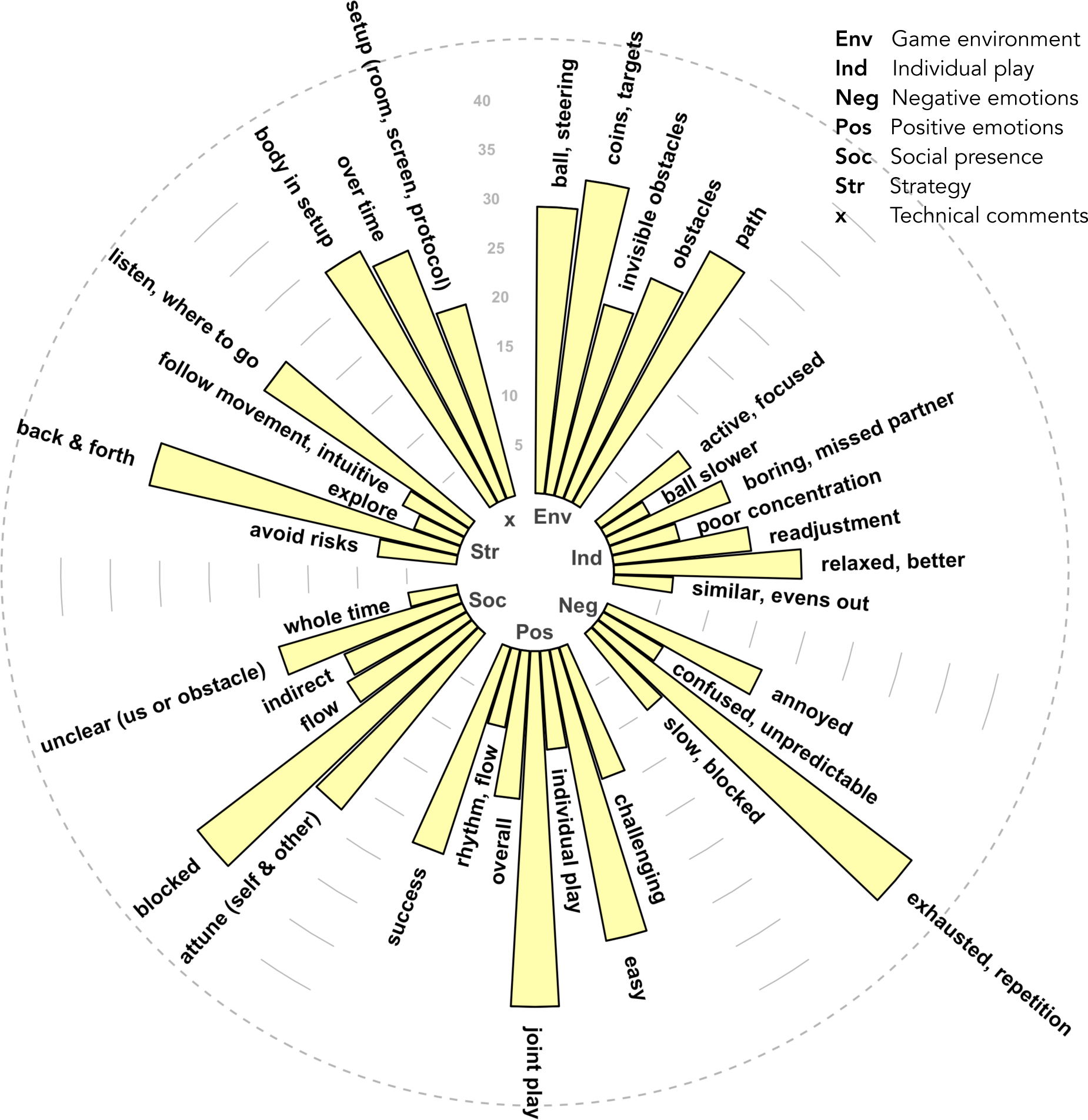
Thematic content of participant interviews. The length of each bar indicates the number of participants that voiced a given code. Grouped bars belong to the same theme (see legend of themes in upper right corner of this figure).

### Predictors of social experience: successful coordination and interpersonal variability

To integrate our observations of the social interaction dynamics in the BallGame, we calculated linear mixed effects models that assess the influence of parameters from each of our levels of observation on participants’ experience ratings (see ***Figure 2***). **Figure 4** illustrates the final models of participants’ engagement, agreement and predictability ratings, respectively. Our findings demonstrate significant influences from members of each class of observation on participants’ social experience: gaming behaviour (*blue* boxes/ predictors in **Figure 4**), movement coordination (*green* predictors), personality differences (*red* predictors), as well as the larger interaction context (*white* predictors).

**Figure 4.**
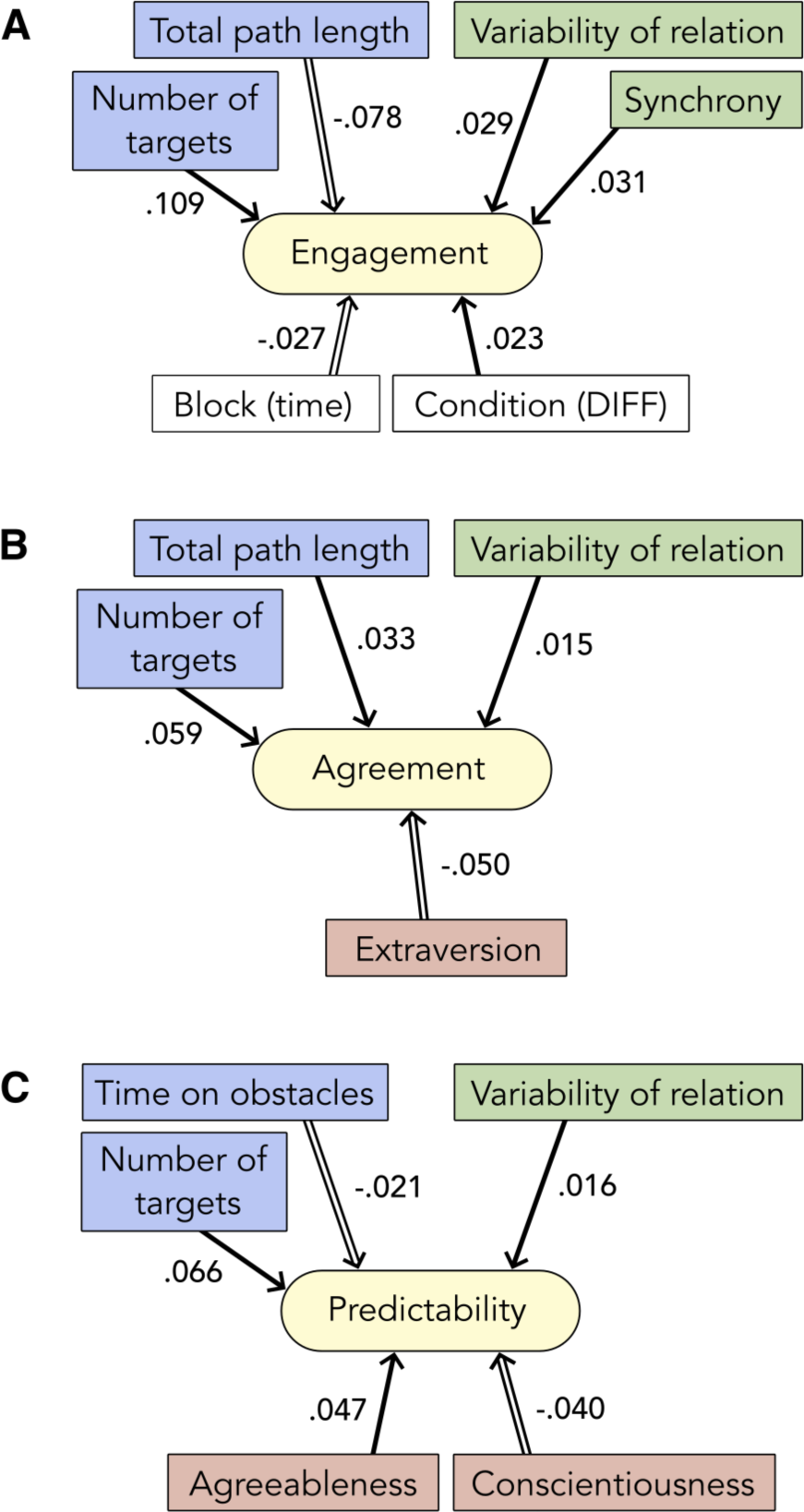
Overview of the final Linear Mixed effects Models of Participants’ Engagement, Agreement and Predictability Ratings (***A*** *to **C***). ***Filled arrows*** indicate positive relations (e.g., longer paths predict higher agreement ratings), ***empty arrows*** indicate negative relations (e.g. later time predicts lower engagement ratings). In line with the overview presented in Figure 2, the colour of the boxes/predictors indicates their class of observation: *(**blue**)* gaming behaviour, *(**green**)* finger movement, *(**red**)* personality traits. Time and game condition, as generic contextual factors, are shown in *(**white**)*. Annotated numbers represent predictor estimates.

Most prominently, we consistently found a higher number of targets collected as well as greater variability in the strength of relation between participants’ finger movements to be associated with enhanced social experience. This is true for the model of participants’ engagement ratings (targets: t = 3.710, p < .001, estimate = .109; variability of relation: t = 2.397, p = .017, estimate = .029), as well as the models of participants’ agreement (targets: t = 3.704, p < .001, estimate = .059; variability of relation: t = 1.971, p = .050, estimate = .015) and predictability ratings (targets: t = 5.784, p < .001, estimate = .066; variability of relation: t = 2.363, p = .019, estimate = .016). The final model of engagement ratings further included synchrony as a positive predictor (t = 2.178, p = .030, estimate = .031), an indicator of general alignment between participants’ steering directions across interpersonal lags. Path length emerged as a significant predictor of both engagement and agreement ratings - negative in the former (longer paths predict lower engagement ratings; t = -3.737, p < .001, estimate = -.078), and positive in the latter case (longer paths predict higher agreement ratings; t = 2.532, p = .012, estimate = .033). Hence, while travelling long distances together stimulated a sense of agreement with one’s partner, it dampened engagement. Relatedly, we found that the further the time in the experiment had progressed, the lower participants rated their engagement (t = -7.260, p < .001, estimate = -.028), in spite of simultaneous improvements in performance and movement coordination over time (see above, *learning effects*, and *Figure SE.1*). Both effects are likely related to fatigue due to repetition or boredom, as participants unanimously reported in the interviews (*Figure 3,* theme ‘*negative emotion*’, sub-category ‘exhausted, repetition’, n = 38 participants). On the other hand, the model of engagement ratings included the joint play condition as a significant predictor: engagement ratings were higher after joint play DIFF trials (t = -2.213, p = .028, estimate = -.023). The increase in engagement in joint play DIFF suggests a stimulating effect of coordinating with a partner that holds a complementary view of one’s environment.

### Additional predictors: obstacle collision and personality differences

The final model of predictability ratings also included obstacle time as a significant negative predictor (t = -2.088, p = .038, estimate = -.021): spending more time on obstacle regions reduced participants’ sense of predictability. We relate this finding to statements in the interviews about having to figure out whether the slowdown was caused by the partner or an invisible obstacle. In this sense, more time on obstacles meant greater potential for confusion, or else, a lack of orientation and predictability.

The final models of agreement and predictability ratings included effects of personality difference (see red boxes in *Figure 4B and 4C*). The differences in conscientiousness between players decreased predictability ratings (t = -2.087, p = .049, estimate = -.040). The association of smaller differences in conscientiousness with higher interpersonal predictability might suggest that similar levels of ambition and discipline make players predictable to each other in this kind of social interaction. Relatedly, we found that greater similarity in trait extraversion - the tendency to be active, optimistic, interested in communication and exciting stimulation - led to higher agreement ratings (t = -2.194, p = .041, estimate = -.050). Accordingly, players may have differed in their tendency to steer the ball through or around obstacles, explore new or repeat old paths, and displayed further more fine-grained differences in steering behaviour. All of these divergences could have caused difficulty to move the ball in a coordinated fashion and thus made it more likely for players to disagree, be stuck on obstacles, and find each other unpredictable. However, we did not find a statistically significant relationship between these personality traits and target sequence complexity (as an indicator of the tendency to explore versus exploit).

Opposite to the effects we observed for extraversion and conscientiousness, differences in trait agreeableness benefited social experience: teams of players with different tendencies for altruistic, empathic, understanding or benevolent behaviour gave higher predictability ratings (t = 2.424, p = .024, estimate = .047). When plotting agreeableness differences against interpersonal time lag, we further found evidence for a positive relation between these two characteristics at small to intermediate levels of agreeableness differences (see *Supplementary Figure SC.2*). This is congruent with the development of more prominent leader-follower relations in pairs with moderate differences in agreeableness.

*Supplementary Materials C* present a complete overview of the initial and final model parameters, as well as the leave-one-out cross-validation that we performed to assess our final models’ generalisability. Predicted and observed ratings correlated significantly in all three cross-validations, speaking to the generalisability of our findings.

### Learning effects: changes in experience and behaviour over blocks and sessions

To assess changes in experience, behaviour and movement coordination over shorter (blocks of 3-4 minutes duration) and longer time intervals (sessions of 20 minutes), we calculated an ANOVA. We found that participants’ performance improved over both blocks (targets: ATS = 8.43, p = .003, obstacle time: ATS = 9.078, p = .005, and path length: ATS = 13.911, p < .001) and sessions (targets: ATS = 29.146, p < .001, obstacle time: ATS = 7.17, p = .024, path length: ATS = 10.296, p = .008), that is, over both short (one block = 3-4 minutes) and intermediate periods of time (one session = 20 minutes). We also saw changes in our measures of movement coordination: mutual information (MI) increased over blocks (ATS = 13.027, p < .001), and both synchrony and strength of relation increased from the first to the second session (synchrony: ATS = 18.647, p < .001, strength of relation: ATS = 20.249, p < .001). *Supplementary Figure E1* visualises these effects, *Supplementary Materials D* provide an overview of statistics, including post-hoc comparisons between individual blocks.

### Differences between the two joint play conditions: effects of seeing the same versus different obstacles

In our ANOVAs we also compared observations across the two joint play conditions, i.e. at the times when participants saw exactly the same or partially different obstacles.

When participants saw the same obstacles, they collected more targets (ATS = 6.807, p = .030), but spent more time on obstacle regions (ATS = 5.896, p = .046). We also found differences in measures of movement coordination between the two joint play conditions: synchrony (ATS = 34.137, p < .001), strength of relation (ATS = 19.913, p < .001) and MI (ATS = 40.549, p < .001) are all higher in joint play SAME. *Supplementary Figure SE.2* visualises these effects of condition for both performance and interpersonal movement coordination measures. Thus, while performance is balanced across the two joint play conditions, movement coordination is higher in joint play SAME.

When considering changes over time, we further saw that MI evolved differently over time under the two joint play conditions, with significant interaction effects of the joint play condition with both block and session, respectively: MI increased over blocks in joint play SAME, but not DIFF (ATS = 9.912, p = .020; see *Figure 5A*), and MI increased more strongly in joint play DIFF, versus SAME, over sessions (ATS = 7.77, p = .046; see *Figure 5B*). Relatedly, we saw that target sequence complexity tended to increase over blocks in joint play DIFF, with an opposite trend in joint play SAME (ATS = 3.75, p = .079; see *Supplementary Figure SE.4D*).

These results provide evidence for slower learning in conditions of different obstacle visibility between players. Having different obstacle visibility also led to higher engagement ratings (see above, Figure 4A). Overall, our findings therefore suggest that a complementary view supports the development of more complex and involving behavioural and coordination dynamics.

**Figure 5.**
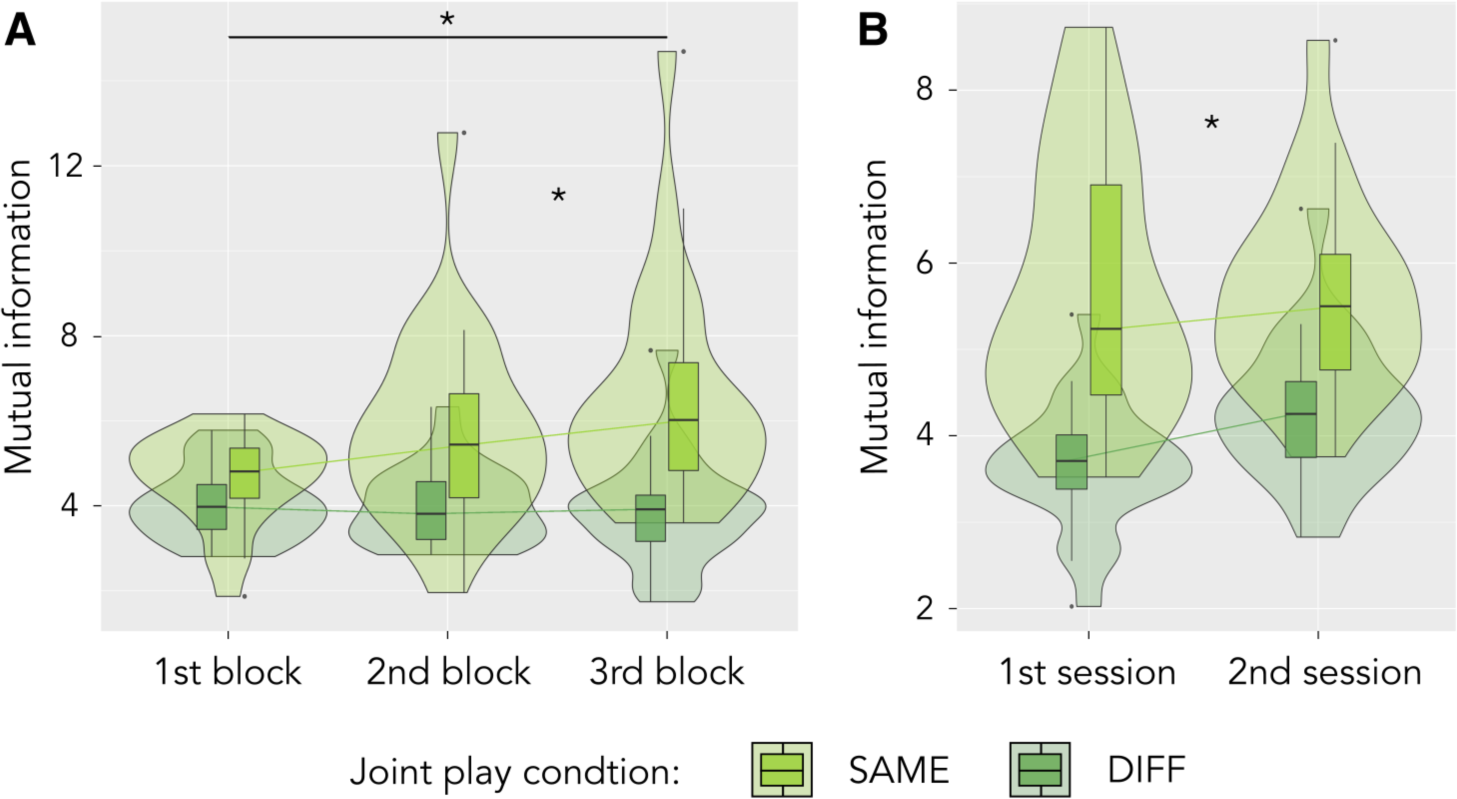
Mutual Information (MI) evolves differently in periods when players see the same (joint play SAME) versus different obstacles (joint play DIFF). Both plots display significant interaction effects as revealed by an ANOVA. Significance levels (FDR-corrected) are indicated by * (p < .05) and *n.s.* (p >= .05). **(A)** Interaction effect of condition and block: MI increased more strongly in joint play SAME from the first and second to the third block. **(B)** Interaction effect of condition and session: MI increased more strongly in joint play DIFF over sessions.

### Individual differences and within-trial dynamics: following up on participant reports

*Individual differences at the transition from joint to individual play.* When asked what it was like to play alone again after the joint play period, participants’ comments ranged from clearly negative (‘boring’, ‘just working it off’, ‘missed my partner’) to rather positive (‘I felt more active’, ‘now I knew how to control the ball and could just to do my thing’). To follow up on these reports, we calculated an ANOVA of performance in highly versus weakly coordinated players. Our results show that coordinated players overall collected more targets (ATS = 17.378, p < .001), and displayed a drop in performance at the shift from joint to individual play - the opposite was true for weakly coordinated players, whose lower performance increased when they shifted to individual play (ATS = 9.903, p = .002; see *Figure 6A*). When comparing the sense of ball control as rated by participants before and after joint play (ball control was only assessed in periods of individual play), we further found that the sense of ball control increased more strongly for players from weakly coordinated pairs (ATS = 4.423, p = .035; see *Figure 6B*). Importantly, players from strongly coordinated pairs indeed talked more negatively about the final period of individual play (see *Supplementary Table SB.1*).

**Figure 6.**
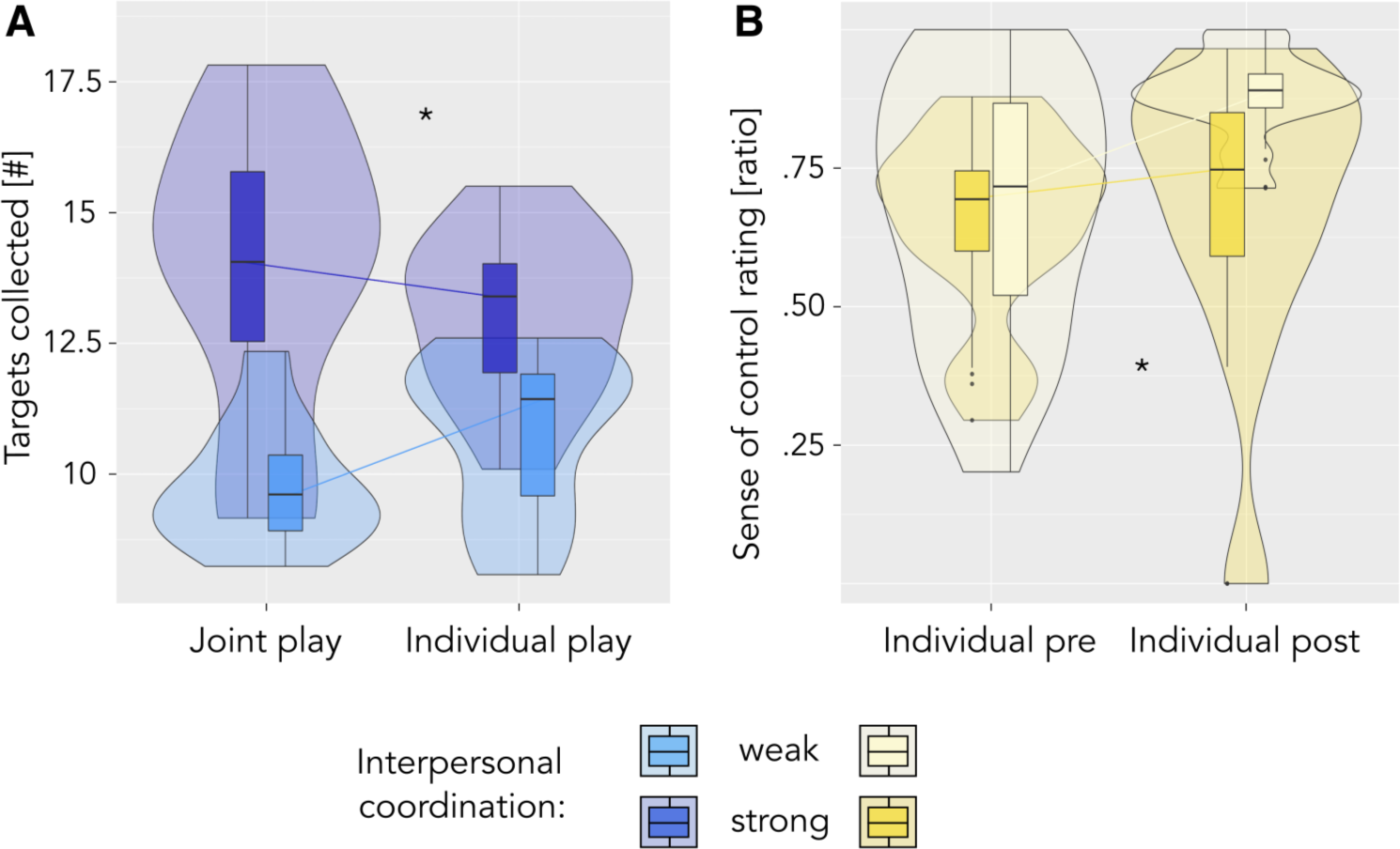
Players from strongly versus weakly coordinated teams at the shift from joint to individual play. Plots display interaction effects of coordination level (strong versus weak) and time. Significant effects are indicated by ** (p < .01) and * (p < .05). **(A)** Players from highly coordinated pairs collect fewer targets in the final period of individual play compared to their joint play performance in the second session - the opposite holds for players from weakly coordinated pairs. **(B)** The sense of ball control increases more strongly from the first to the last 10 trials of individual play for players from weakly coordinated pairs.

*Within-trial changes in the gaming dynamics*. Participants described a marked shift in their experience from the early trial, during which they were deeply involved with resolving coordination issues and understanding what their partner wanted, to the late trial, which was more about performing and could even feel like playing alone. To trace these within-trial changes in game-concurrent observations, we calculated an ANOVA of performance and movement coordination measures within trials, cutting the trial into three segments of 20 seconds each. Our results show that coordination improved significantly and rather continuously throughout the trial (see *Figure 7A to C;* synchrony: p = .006, ATS = 8.51; strength of relation: p < .001, ATS = 13.26; time lag: p = .006, ATS = 8.48). Performance measures, however, evolved in a less regular fashion: while generally improving over trial segments (see *Figure 7D to F;* targets: p < .001, ATS = 19.73; obstacle time: p < .001, ATS = 155; path length: p < .001, ATS = 102.81), the number of targets collected as well as the total path length increased mostly from the first to the second trial segment, whereas obstacle time only dropped in the final trial-third (see *Table SD.3* for exact post-hoc pair-wise comparison statistics). These findings agree with participant statements about changes in the gaming dynamics across the trial. They also indicate that participants succeeded relatively quickly at reaching more targets, whereas reducing obstacle collisions was only accomplished late during the trial.

**Figure 7.**
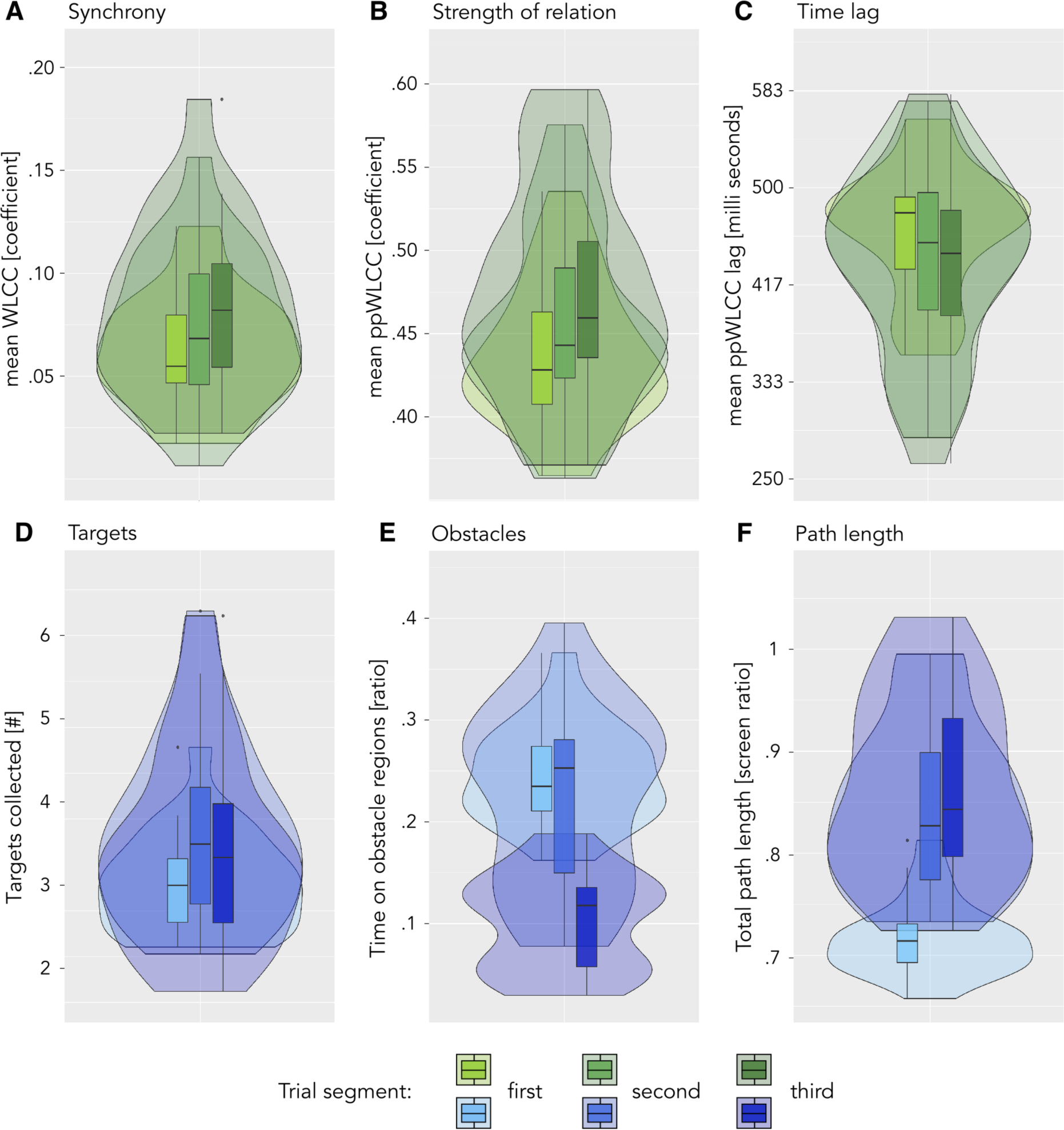
Within-trial changes in movement coordination and gaming behavioural measures. Plots display significant main effects of trial segment as revealed by ANOVAs. Significance levels (within-class FDR-corrected) are indicated by *** (p < .001), ** (p < .01) and * (p < .05). Top row, significant effects in movement coordination measures: **(A and B)** Synchrony and Strength of Relation increased from the first and second to the final trial segment. **(C)** Interpersonal time-lag was lower in the second and third, compared to the first trial segment. Bottom row, significant effects in performance measures: **(D)** The number of targets collected was lowest in the first and highest in the second trial segment, after which it decreased again from the second to the third trial segment. **(E)** Obstacle time was only reduced in the third, compared to the first and second trial segments. **(F)** Path length increased from the first to the second, as well as from second to third trial segment. See tables SD.4 and SD.3 for an overview of statistics.

*Game objects as attractors of attention.* Participants frequently mentioned all objects present in the game environment - to recall their experience, describe strategies used in the game or voice emotions. This prompted us to investigate coordination in the vicinity of targets and obstacles. More specifically, we compared the strength of relation between players’ steering movements across the target cycle, as a function of how long ago the last target was collected and how quickly they would reach the next one. As can be seen in *Figure 8A*, the strength of relation between participants’ steering movements was highest during and immediately after target collection. This pattern was remarkably consistent across both joint play conditions (plotted in light versus dark green, *Figure 8A*), with a significant, consistent offset in the strength of relation between the two conditions. We also compared the strength of relation at times when the ball was closest to an obstacle that both or at most one player could see: players’ steering movements were more strongly related when the ball was closest to an obstacle that both of them could see (ATS = 57.247, p < .001; see *Figure 8B*). In short, objects in the game environment showed to have a strong influence on interpersonal coordination.

**Figure 8.**
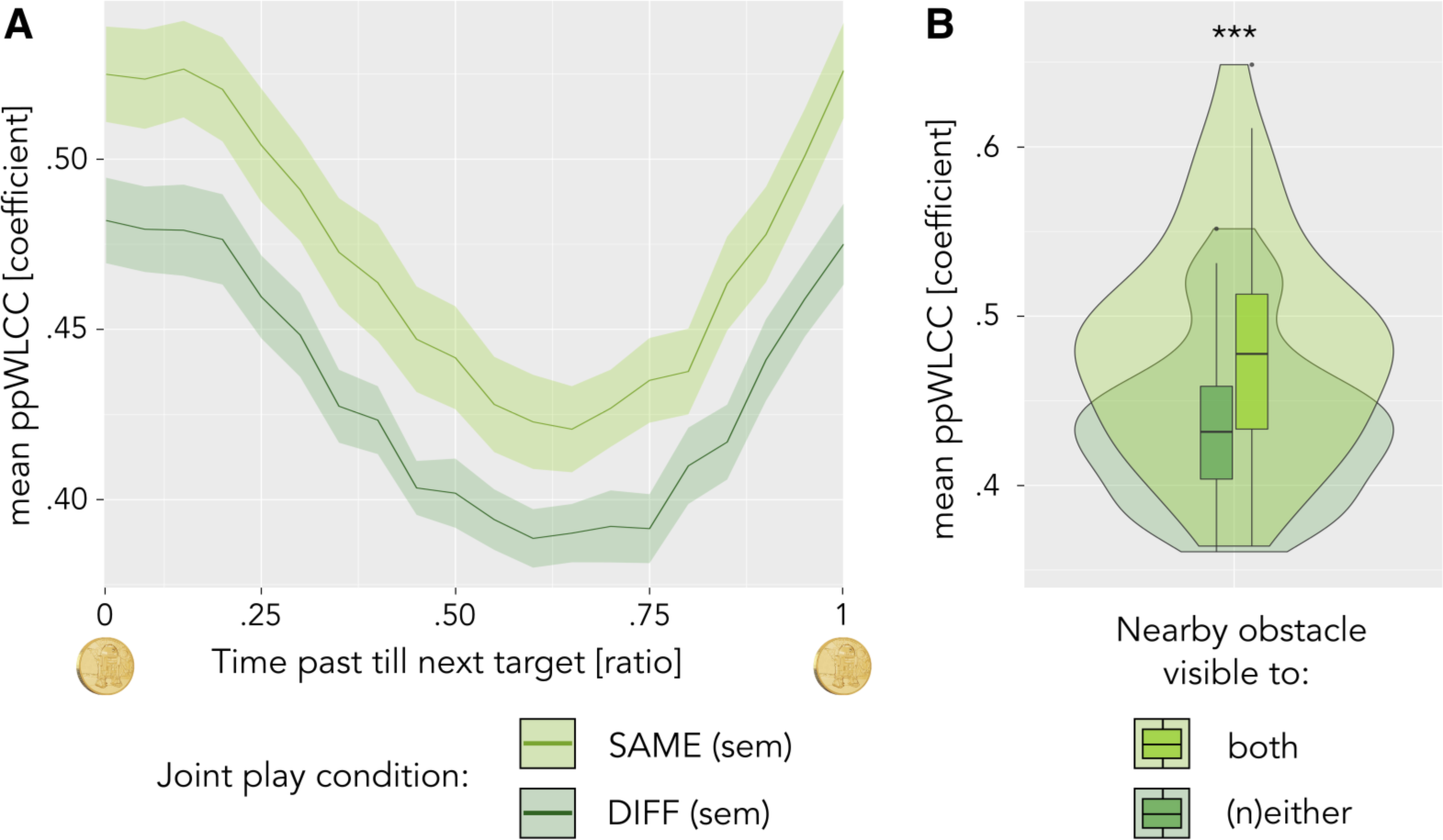
Objects in the game environment influence interpersonal movement coordination. FDR-corrected significance levels are indicated by *** (p < .001). **(A)** Coordination across the target cycle: lines and shaded areas represent mean and standard error of the mean strength of relation between two target collection events in the two joint play conditions (joint play SAME = dark green, joint play DIFF = light green). **(B)** Coordination as a function of obstacle visibility: players’ steering movements are more strongly related when the ball is closest to an obstacle that both players can see (light green bar and violin plot), versus neither or only one of them (dark green bar and violin plot).

## Discussion

The present work contributes to social interaction research in three distinct ways. First, we present a novel paradigm for the study of motivated and continuous interaction. Second, our analyses integrated multiple levels of observation, including in particular several operationalisations of sensorimotor coupling in a social context. Besides highlighting moments of target collection and variability in the strength of interpersonal coordination as predictors of enhanced social experience, this allowed us to describe the influence of movement coordination, gaming behaviour, personality traits and the interaction context on social experience. Third, following participant reports, we identified the degree of coordination as a marker of individual differences in experience at the transition from joint back to individual play, were able to locate the evolution of interpersonal coordination dynamics in the game environment, and revealed marked and divergent changes in gaming behaviour and movement coordination over short periods of time (within the trial). Overall, our findings emphasise the specific temporal and spatial contexts for interpersonal coordination, and point to a critical balance between interpersonal synchrony and difference for positive social engagement.

### Social engagement requires interpersonal synchrony and variability

Our findings strongly suggest that both ‘synchrony’ (e.g. strength of relation, a shared view, collecting targets) and ‘difference’ (e.g. variability of relation, a complementary view) are important for positive social experiences. Good performance and synchrony are straightforward predictors of experience - both contribute directly to the task at hand (‘coordinate with your partner and collect as many targets as possible’). An increase in the variability of the peak strength of relation between players’ steering movements (independent of the time-lag between them) is a more surprising predictor of experience. It concurs with the opposite effects of path length on engagement and agreement, the decrease in engagement over time and its increase after joint play DIFF trials, in indicating an important balance between predictability (successful coordination) and stimulating difference (fun, challenge, surprise) for engaging social interactions. A special role for variability in social interactions - next to a positive impact of synchronous or otherwise coordinated behaviour - is in line with Proksch and colleagues’ (2022) finding of a parallel increase in stability *and* variability over the course of an orchestra performance explicitly designed to transition from uncoordinated to coordinated behaviour. The authors applied recurrence quantification analysis to sound recordings of the performance - variability in this case referred to recurrent sound (amplitude) sequences of variable length. Early work on interactional synchrony in the movements and vocalisations of a conversing group further points out variations in interpersonal synchrony as a key mechanism for coordinating switches in communicative roles (Kendon, 1970). Importantly, studying the creativity of joint productions as a function of the groups’ conversational style (instructive vs. inclusive vs. integrative), Bjørndahl and colleagues (2015) found that creativity is high when group members synthesise (integrate) different viewpoints (by engaging in frequent repair of own and others’ contributions). We also find support for a contribution of both alignment and difference in the theoretical literature. De Jaegher’s (2007) account of cognition as participatory sense making, for example, puts variability centre stage: “social interactional timing is a variable affair, and not rigid”. Similar to the suggested balance between synchrony and difference, authors from diverse fields have emphasised the importance of both exploiting proven and exploring new strategies: from healthy psychological attachment (Bowlby, 1982), to successful foraging behaviour (Stephens et al., 2008; Hills et al., 2015), creativity (Hart et al., 2017), persistence and having fun (McCullough, 2013). The suggested balance between synchrony and difference can also be compared to the co-existence of integrative and segregative tendencies that is emphasised in dynamical systems theory (Tognoli & Kelso, 2014). Overall, we find strong evidence for an important role of variation in social sensorimotor coordination: successful interpersonal engagement seems to require room and sensitivity for differences just as much as it relies on the capacity to integrate them in synchronised forms of acting and shared understanding (see also Sebanz et al., 2006).

### Rhythms of coordination: invariant features in the environment as guidelines for interpersonal attention

Participant reports and our follow-up analyses revealed strong local modulations of interpersonal coordination and experience in space (around targets and obstacles) and time (within the trial and at the transition from joint back to individual play). In particular, our findings suggest several scales of recurrence through which the link between external structures and ongoing interpersonal coordination is established and maintained. The notion of several time-scales of entrainment is in line with the distinction between modality- and object-specific sensorimotor contingencies (O’Regan & Noë, 2001; see also Maye & Engel, 2012). Applied to the social domain, this implies that engaged coordination requires participants to align their attention through shared features in the environment: from immediate sensory (e.g. approaching a target or obstacle, steering a particular curve) to longer-term frames of reference (e.g. typical course of a trial, gaming behavioural strategies, roles in the interaction). Taking a very similar approach, predictive coding theories of neural and cognitive function (e.g. Clark, 2016) suggest that cognitive functions (and our nervous system) continuously generate schemas and models about meaningful aspects of our environment (predictions, hypotheses), and engage in exploratory behaviour that probes, fine-tunes, extends or repurposes them. In successful social interactions, this balancing between reliability and effective exploration becomes a shared process. Importantly, Jones’ dynamic attending theory (1976, 2019) and Lefebvre’s rhythm analysis (2004) teach us that rhythmic coordination is at the same time recurrent and *progressively changing*. Thus, besides emphasising local (situated, individually different) histories of coordination, a rhythmic perspective suggests the investigation of coordination from one instance to the next (in our case e.g. the evolution of coordination between subsequent target collections, obstacle collisions, trials, blocks, etc.).

### A call for future work on the particular kind of dynamics that sustain interpersonal coordination

Our approach does not yield simple (nor final) answers about the relationship between interactive movement coordination and social experience.

First, the analysis and report of the game-concurrent eye-tracking and EEG recordings is beyond the scope of this article. However, these data present a well-suited extension to answer our research questions on a neurophysiological level. For example, in light of previous findings that show enhanced movement driven modulation of neural activity in followers (especially Zhou et al., 2016, who observed repetitive hand opening and closing; see also Dumas et al., 2010, who studied spontaneous imitation of hand movements), it would be interesting to assess whether ‘follower-typical’ neural modulation is enhanced in participants with higher levels of agreeableness. Likewise, one of the planned next analysis steps is to look for neural correlates of the experiential and behavioural changes we observed within the trial, to explore potential substrates of a shift in focus from social to performative. Beyond that, the consideration of eye movement, pupil dilation and EEG data is likely to complement our understanding of social interaction in the BallGame.

The BallGame presents an example of combining quantitative and qualitative methods to trace continuous interaction dynamics. Future research is needed to refine such multi-level approaches to studying social engagement. Efforts into this direction can, for example, be found in Hall and Stevens’ (2015) Interaction Analysis, a method for reconstructing gestural and conversational interactions in a group of people. Similarly, the investigation of free play in groups of children as presented by Kalaydjian and colleagues (2022) highlights gestures of suggestion, recognition and confirmation as different phases of a joint (distributed) transition between making, following and breaking rules. Incorporating both momentary and aggregated measures of behaviour and experience, as well as paying attention to how coordination unfolds through recurrent, progressively changing patterns that attune to the local context appears to be key. This implies that future work should also deliver extended modelling approaches, for example to estimate mediating relations between individual predictors and a greater variety of temporal dependencies.

Our main finding about a positive contribution of variability and difference to engaging social interactions likewise merits greater attention. One strategy for pursuing this could be especially designed experience assessment (such as questions about challenging, creative or surprising moments; see also the dynamic ratings of togetherness in Noy et al., 2015; or the phenomenological investigation of interaction dynamics in Kimmel et al., 2018). Another approach could be to ease the contribution of difference to social interactions in the laboratory. This could be accomplished by measures such as starting the experiment with a simple activity that allows participants to notice and express their experience, or by using an experimental task that provides a clear frame but invites creative contribution.

### The BallGame as a novel paradigm in social interaction research

The BallGame can be located between tasks that require rhythmic interpersonal coordination (Konvalinka et al., 2010; Dumas et al., 2010; Dumas et al., 2014a; Llobera et al., 2016; Zhou et al., 2016; Vesper et al., 2016; Varlet et al., 2020) and experimental approaches that focus on natural interactions (Ramseyer & Tschacher, 2016; Kimmel et al., 2018; Jakubowski et al., 2020). Our design is explicitly set up at their intersection: interactional synchrony provides an advantage but is not the only or explicit goal in the BallGame. Communicating and finding agreement based on individual preferences and complementary viewpoints and actions is equally relevant. Importantly, the BallGame involves continuous interpersonal sensing and acting, overlapping possibilities for action and shared as well as complementary information between players. As such, this task encourages social engagement and leaves room for individual and interactional autonomy around the development of interpersonal coordination. Such an approach entails analytical challenges: greater freedom implies greater potential for genuine social engagement, but also more complex interpersonal dynamics that are harder to capture in simple measures of behaviour and interpersonal coordination. Therefore, we have chosen an iterative approach that goes beyond generating detailed multi-level records and assessing changes over time: we followed up on participants’ specific descriptions of their interpersonal experience. We argue that this combination provides a powerful tool and a novel approach in laboratory studies of social learning and engagement. In conclusion, we urge future studies to leave room for participant autonomy and invest in thorough evaluation of commonalities and differences in their experience. Analytic approaches can then be designed to integrate what is learnt about the specific histories of coordination that unfold between participants.

## Acknowledgements

This work was supported by grants from the EU (project ‘socSMCs,’ H2020-641321) and the DFG (SFB936-178316478-A3 and TRR169-261402652-B1/B4). Karin Deazle provided invaluable support with participant recruitment and preparation. Hanna Krause contributed to the development of the semi-structured interview.

## Author contributions

AL and FG developed the experimental paradigm and analytic strategy in exchange with TS and AKE. MS, AL and FG implemented the hard- and software of the game environment and hyper scanning setup. AL and FG conducted the study and analysed the data in exchange with TS. AL and KH worked on the qualitative analysis. AL wrote the article in close collaboration with FG. AKE acquired the funding and supervised the study. All authors reviewed the manuscript.

## Data availability statement

Data and code are available upon request.

## Additional information

The authors declare no competing interests.

