## Supplementary Materials for "Predicting social experience from dyadic interaction dynamics: the BallGame, a novel paradigm to study social engagement"

*A. Movement coordination measures*
  *1. Detailed methods*
    *I. Windowed-lagged cross-correlation (WLCC)*
    *II. Mutual information (MI)*
    *III. Phase-slope-index (PSI)*
  *2. WLCC Surrogate statistics: real vs surrogate correlation across interpersonal lags*
*B. The Interview*
  *1. Protocol*
  *2. Thematic content analysis*
*C. Linear Mixed Effects Models*
  *1. Initial and final model parameters*
  *2. Cross-validation of the final models' fixed effects*
  *3. Plotting agreeableness differences against interpersonal time lag*
*D. MANOVAs and post hoc ANOVA statistics, including pair-wise comparisons between individual blocks and trial segments*
*E. Additional visualisations of ANOVA results*
  *1. Main effects of block, session and condition*
  *2. Main effects of trial segment (basic movement)*
  *3. Additional interaction effects (trends)*

### Supplementary Materials A

### Movement coordination measures

#### 1. Detailed methods

***Windowed-lagged cross-correlation****:* In line with a recent validation study (Schoenherr et al, 2019), as well as our own tests with a range of parameter settings, we settled on windows of 3.6 seconds (220 frames) and a window overlap of 3.3 seconds (200 frames), leading to a step-size of 0.3 seconds. We further assessed correlation between players' movements at interpersonal lags of up to 1.1 seconds (70 frames) with single frame increments (60 Hz resolution, i.e. 0.017 seconds). In other words, we assessed correlation across 141 lags (70 lags player one leads, 70 lags player two leads, 1 lag simultaneous movement) at 157 time points during each trial. In other words, we assume that events last approximately 3.6 seconds, that the evolution of players' interactions can be observed at a resolution of 0.3 seconds, and that inter-player coordination does not exceed a delay of 1.1 seconds and can be observed at a resolution of 0.017 seconds.

***Peak-picking***: To identify potential leading or lagging between players at each of the 157 time steps, we identified the lag between players' steering directions that showed the strongest correlation above a minimum of 0.3 that was closest to simultaneous movement (0 lag). Since we set time steps at which the maximal correlation coefficient over all lags is lower than 0.3 to zero, our ppWLCC parameters indicate coordination over short, intermittent periods of time

***Mutual Information***: MI is a framework that allows for the quantification of shared information between two signals and is based on Shannon's entropy from information theory. Entropy is computed by binning the given data set, and calculating the probability of a given data point to fall into either of these bins. These probability values are then multiplied with the logarithm of the probabilities, summed and multiplied by minus one to return to a positive scale. MI is calculated by adding the individual entropies of the two signals and then subtracting their joint entropy (Cohen, 2014). Using custom-made scripts based on Cohen (ibid), we calculated MI values for each block, that is, in parallel with moments of experience rating in the experiment. A block comprises three or four trials. Accordingly, we calculated MI for durations of 180 or 240 seconds, as well as for each session, condition and pair. The number of bins to discretise the data was estimated using the Freedman-Diaconis rule (Freedman & Diaconis, 1981). As the number of bins influences the entropy estimate, we first estimated the optimal bin number for each pair, session, condition, and block. Then, we took the ceiling of the grand average of the optimal bin number (25 in our data set) and re-ran the whole analysis with 25 bins for every calculation. In order to derive standard statistical Z-values of our MI estimates, we additionally applied permutation statistics. In 500 iterations, we temporally shifted one of the two time series by some random factor and calculated MI, to finally create a distribution of MI values expected under the null hypothesis from which Z-values were derived (again compare Cohen, 2014). These Z-values (one for every pair, session, condition, and block) were then subjected to statistical comparisons.

***Phase-slope-index***: As for MI, we aggregated data at the level of the block. We estimated PSI (using data2psi.m, METH toolbox, Guido Nolte) for segment lengths ranging from 20 samples (0.333 seconds at a sampling rate of 60 Hz) to 240 samples (4 seconds) in steps of 20 samples (0.333 seconds) but for further analysis, we selected PSI values calculated using a time window of 40 samples (0.667 seconds). This choice was motivated by systematical assessment of the parameter space in an initial analysis step with the goal of finding the segment length that maximises the mean normalised PSI (calculation of the standard deviation across epochs using the jackknife method was done with different epoch lengths of 2, 3, and 4 seconds) across pairs, sessions, conditions, and blocks. What is more, PSI was calculated across all frequencies (wide band) based upon visual inspection of the grand average power spectra, where no apparent frequency peaks nor differences between conditions were present.

### 2. WLCC Surrogate statistics: real vs surrogate correlation across interpersonal lags

***Surrogate calculation****:* We calculated surrogate WLCC on data of players from distinct pairs moving through the same game landscape. To take into account the potential influence of both temporal and spatial context, we preserved the temporal structure of the trial as well as the spatial arrangement of targets and obstacles. Figure S.1 illustrates the mean (and ci of the mean) WLCC for real compared to surrogate pairs across the interpersonal lags we considered in the remainder of our analyses. Both statistics are based on mean values across all pairs and blocks of joint play.

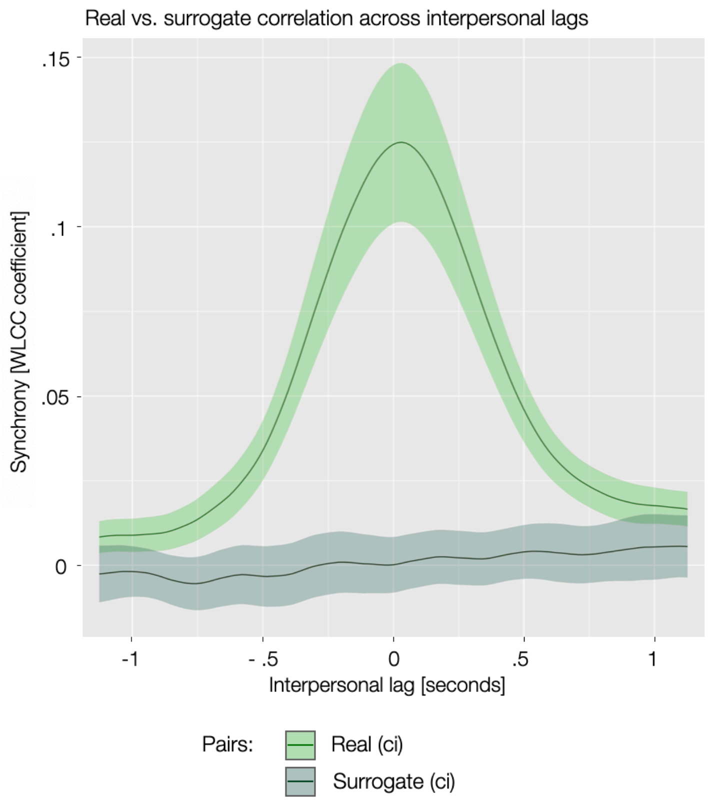


**Figure SA.1** Synchrony in real and surrogate pairs. Mean windowed-lagged cross-correlation coefficients are displayed for real and surrogate pairs separately across interpersonal lags of up to 1.27 seconds. The shaded area indicates the confidence interval (ci) around the mean.

**Supplementary Materials B**

### The Interview

#### 1. Protocol

*Open questions:*

*Q1 - When you think back to playing the BallGame, what comes to mind?*
*Q2 - During the BallGame, what was most present for you, what did you focus on?*
*Q3 - Is there anything that was (a) easy, (b) exhausting or (c) fun?*
*Q4 - What would you compare your experience of the BallGame to? (games, everyday life) **
*Q5 - Was your partner present to you? If so, when and how?*
*Q6 - What was it like to play alone again in the end, after the joint play period?*
*Q7 - Did you play according to one or several specific strategies?*
*Q8 - Is there anything you would like to add?*

*Scaled questions:*

*Q9 - How easy or difficult was it to control the movement of the ball through your finger movements? **
- *very easy 0 to 10 very difficult*
- *individual vs joint play*
- *early vs late in the game*

*Q10 - Did the controller irritate you? **
- *not at all 0 to 10 very much so*
- *early vs late in the game*

*Q11 - Was it easy for you to learn and orient within the BallGame? **
- *very easy 0 to 10 very difficult*
- *individual vs joint play*

*Q12 - How did you experience joint play in the BallGame: more as a computer task, or more as a social interaction?*
- *PC 0 to 10 social*

*Q13 - What characterises social interactions that you (do not) enjoy? **
*Q14 - (When) Do you enjoy leading, or following another person's leadership? **

** Based on redundancy (with other questions) and/or irrelevance (to the specific research questions we focused on), answers to these questions were not included in the present analysis.*

#### 2. Thematic content analysis

Most of the themes identified in our thematic content analysis reflect participants' answers to one of our interview questions. For instance, we asked participants what it was like to play alone after the joint play period, and one of the themes that emerged from our analysis was 'Individual play'. Interestingly, we neither asked participants about the game environment, nor the technical setup, themes that nearly all players spoke about both in the early, openly phrased questions, as well as at later stages of the interview. Below, we summarise participant statements in each theme.

**Game environment**. The visual appearance of the game environment was present in detail to participants during the interview. This is observable from remarks such as "there was R2D2 on the coins..", or "the red bars come to mind..". Participants associated targets with a sense of reward, were eager to find a good path and avoid obstacles, including remembering the location of invisible obstacles. For instance, many players reportedly sought to "find the shortest / easiest path!", "go fast from A to B" and often recalled "the obstacles.. being slowed down" or, "especially when together - where are the invisible obstacles?". Participants also used the game environment to describe strategic considerations - "this moment, before each trial - we could think about which path to take"; or, "it's better go in curves, the direct path is usually blocked". Participants further mention their sense of (not) being able to reach coins, steer the ball faster, avoid obstacles or get off them quickly. For instance, many players expressed that "it was complicated - is there an invisible obstacle? Are we in disagreement? This not-knowing..". Another player pointed out that it was "frustrating to be stopped right before the coins, even more when playing together". On the other hand, many players described the pleasure of "that moment of - ah, here is a path!".

**Positive emotion**. Participants clearly expressed their overall positive experience of the game: "it was fun! I didn't check the time"; "really cool experiment! No 'experiment experience' for me!"; "I was deeply engaged"; "I was motivated to collect the coins - you really enter the game!". They also referred to the rewardingly challenging nature of the task - "it's not easy but you can figure it out! many small moments of success - difficult, but, we made it: I am steering by myself! that's how it felt!". Besides success ("reaching targets"; "when we managed to quickly identify our path"), positive emotions were was associated with joint play - "fun? playing together! not only coins, also coordinating! Learning about the other, how to coordinate"; "being able to steer through small paths between obstacles - nice, he thinks exactly like me!"; "when it worked well together, I had to suppress laughter!"; "being fluent"; "on one wavelength"; "together and unanimous - nice!"; "over time, more control, new paths, more fluent"; or "after a while, having found a rhythm, coordinating just through our finger-movements, cool". Another group of positive statements was expressed around ease: "the steering was easier than I thought!"; "moving the ball, easy, like an extension of the body"; "when you had a feeling for what the other wants, then it was easy / good"; "the back and forth strategy - I was relieved to see that we easily settled on that". Another group of positive statements was made about individual play: "fun? playing alone. when together, there were irritations"; or, "alone much more fun, invisible obstacles easy.. yes..".

**Negative emotion**. Participants described sensations of exhaustion and frustration. They frequently mentioned "these annoying invisible obstacles", and complained ("come on…!") about coordination difficulties - "when the ball goes into another direction, not the one I wanted"; "when you tried, but you were stopped"; "obstacles, being blocked, slowed down, not being able to have my way - that was frustrating"; "when the other sees the obstacle, but cannot simply decide where we go - when it took longer than necessary to solve a block/obstacle". Confusion or lack of clarity was another source of negative mood: "somehow it was a bit stressful, not knowing what the other wants now"; "steering was somehow strange, as if it didn't always react the same way"; "exhausting. I felt like I couldn't do much - I tried to listen and respond to her strategy - sometimes it worked, sometimes it didn't". Participants further reported boredom and tiredness during the late part of the game: "exhausting? The duration / repetition"; "the coins appeared at predictable spots, that could be exploited (back and forth strategy / monotony)"; "even though I had mastered the finger-thing, it was more exhausting in the end, I simply couldn't focus, tired", or, "the control tasks were really exhausting, just staring at the screen".

**Social presence.** Particularly the early trial was described as social: "I focused, consciously, especially in the beginning of a trial: where does my partner want to go?"; or, "when playing together - waiting, what does my partner want - is what I want against this? once this is settled, it is like playing alone". In general, participants emphasised mutual consideration and described a process of attuning and coming to agreement: "more just coordinating, coming to agreement..";

“working as a team, trying to understand the intentions of the other”; and “coordination, thinking of one’s partner, being considerate - brings us faster to our goal!”. There were also remarks about flow and fluency: “I had a good feeling for my partner - she didn’t block me and we could always find a path together”; “when it was clear what we wanted to do, then my partner was especially present to me”. On the other hand, many participants report a sense of social presence in moments of disrupted flow: “thinking of my partner? When blocked..”. Relatedly, a group of comments evolved around confusion and the indirect nature of communicating: “when it didn’t go well - why? What does she want? Does she have an obstacle? Why can’t we move on”; “in part funny - you don’t know if it’s an invisible obstacle or your partner who doesn’t collaborate”; “not knowing when you see different things..”; and “not knowing what the other is doing..”, hence “you make an effort, try to communicate, signal..” - “steering, collecting coins with someone you can’t see, no verbal back and forth”. For some, their partner felt present the whole time - “when it didn’t work, or, when it went really well - increasingly, over the course of the game”; “I had a ‘we-feeling’, rather than thinking of her explicitly..”; “throughout the game - where does he go, what does he want, I want him to go here now, or, argh, he wants something else.. everything really!”.

**Strategy**. Participants talked about strategies to coordinate as well as reach pragmatic goals. They described “waiting, to know where the partner wants to go, then support her”; “trying to perceive the other, their intentions - especially in the beginning of a trial”. They also sought to keep track of “invisible obstacles - know where they are, so I can let my partner lead around them”, and “notice when partner wants to avoid an obstacle - respond to that, because I cannot see it. Listen”. To avoid risks (“it’s difficult enough to coordinate with the other - better keep it simple!”), many preferred a ‘back-and-forth’ strategy: “find simplest path between two targets, and stay there”. However, some participants wanted to explore, “not just go back and forth”, “especially when alone - try out more”. Finally, some remarks highlight the intuitive nature of finding a workable path: “steering the ball, like, so that it moves - if it doesn’t, better ‘let go’, go with the other”; or, “this direction doesn’t work? Stay calm, make a few moves, see where it wants to go”.

**Individual play.** Many participants voiced a sense of ease (‘relaxed’) at the transition to individual play - it was easier to play the game - “to know where the invisible obstacles are, when playing alone, easy!”; “it was better to play alone - you could control the ball, find and stick to the shortest path”. Some even described a more active sense of engagement: “when I played alone, I felt more active”; “I really know how this works now, the steering, and I can choose”; “I explored more when I played alone”. Other participants reported a need to readjust: “the steering was unusual - my partner must have kept me on track before!”; “steering a really nice smooth curve - that was better together, being ‘fluent’!”; or even felt that the ball moved slower, “alone was slower than together!”; “I felt like I was slower, alone”[1]. Many participants further felt bored or missed their partner: “the steering was ok but it was a bit boring.. together it was more fun!”; “just going through the motions, working it off”; “it was a bit boring in the end - I missed my partner, when I didn’t want to move, she helped me”; “in the beginning of the session, it was fun to play alone (I didn’t yet know how to play) - in the end not anymore, it was boring”. Finally, several participants found joint and individual play more or less the same - “back and forth, if it doesn’t work, swap route - very similar alone and together!”; “not such a big difference, similar performance”, or, “sort of evens out - easier because of the ball control, more difficult because of three completely invisible obstacles”.

**Technical comments**. Nearly all participants commented on the laboratory environment, either speculating about the technical implementation or mentioning their bodily experience in

[1] While acceleration was reduced when playing alone, the maximum speed of the ball was the same.

relation to the setup. For instance, participants were “thinking about how you realised this - two separate rooms, steering one ball together, all the recordings..”; “noticed the symbols in the corners - for the eye-tracker?”; found it “easy to understand - the game mechanism, with the landscapes etc”; or remarked on the “unusual game control!”. They also mentioned their fingers (steering), their back (sitting) and eyes (staring at the screen). Participants also described general shifts in experience over time - “first figuring out, then fun, then repetitive (lack of focus)”; “at first it was difficult to know what she wants, that got better over time - okay, I have to let go sometimes, notice where she wants to go”, as well as, “the steering got very easy, over time”.

| | strong coordination | weak coordination |
|---|---|---|
| negative statements | 22 | 13 |
| positive statements | 19 | 23 |

**Table SB.1** Statements about the final block of individual play by players from strongly versus weakly coordinated pairs. Negative comments about the final period of individual play are made predominantly by players from strongly coordinated pairs (upper row in the table). Weakly coordinated pairs talk positively, rather than negatively, about the final period of individual play (right column in the table). Negative statements subsumes the codes ‘ball slower’, ‘boring, missed partner’, ‘poor concentration’ as well as ‘readjustment’. All other codes in the theme ‘Individual Play’ are counted as positive statements.

**Supplementary Materials C**

### 1. Initial and final model parameters

*Engagement*

| ***Initial*** **Predictor of engagement** | **estimate** | **std. error** | **t value** | **p value** |
|---|---|---|---|---|
| **Intercept** | **1.193** | **0.049** | **24.207** | **0.000** |
| Autoregressive covariance | 0.022 | 0.069 | 0.324 | 0.748 |
| **Block (time)** | **-0.026** | **0.004** | **-6.950** | **0.000** |
| **Condition (same vs diff)** | **-0.025** | **0.012** | **-2.022** | **0.044** |
| Neuroticism differences | 0.036 | 0.042 | 0.856 | 0.401 |
| Extraversion differences | -0.092 | 0.041 | -2.269 | 0.034 |
| Openness differences | 0.002 | 0.040 | 0.049 | 0.961 |
| Agreeableness differences | 0.024 | 0.044 | 0.529 | 0.602 |
| Conscientiousness differences | -0.010 | 0.041 | -0.242 | 0.811 |
| Autism quotient differences | -0.008 | 0.043 | -0.183 | 0.857 |
| Empathy differences | -0.064 | 0.039 | -1.630 | 0.117 |
| Distance covered by finger moves | 0.048 | 0.030 | 1.584 | 0.114 |
| Number of finger moves | -0.019 | 0.022 | -0.860 | 0.391 |
| **Synchrony (WLCC-coef-mean)** | **0.038** | **0.022** | **1.738** | **0.083** |
| Strength of relation (ppWLCC-coef-mean) | -0.027 | 0.028 | -0.974 | 0.331 |
| **Variability of relation (ppWLCC-coef-std)** | **0.037** | **0.012** | **2.959** | **0.003** |
| Time-lag (ppWLCC-lag-mean) | -0.012 | 0.021 | -0.582 | 0.561 |
| Switching (ppWLCC-lag-std) | -0.012 | 0.014 | -0.845 | 0.399 |
| Mutual information | 0.016 | 0.015 | 1.074 | 0.284 |
| Phase-slope index | 0.008 | 0.013 | 0.580 | 0.563 |
| **Number of targets** | **0.094** | **0.033** | **2.826** | **0.005** |
| Target sequence complexity | 0.029 | 0.022 | 1.346 | 0.179 |
| Time on obstacles | 0.004 | 0.032 | 0.123 | 0.902 |
| **Total path length** | **-0.070** | **0.035** | **-2.004** | **0.046** |

| ***Final*** **Predictor of engagement** | **estimate** | **std. error** | **t value** | **p value** |
|---|---|---|---|---|
| Intercept | 1.199 | 0.055 | 21.703 | 0.000 |
| Autoregressive covariance | 0.021 | 0.069 | 0.311 | 0.760 |
| Block (time) | -0.027 | 0.004 | -7.260 | 0.000 |
| Condition (same vs diff) | -0.023 | 0.011 | -2.213 | 0.028 |
| Synchrony | 0.031 | 0.014 | 2.178 | 0.030 |
| Variability of relation | 0.029 | 0.012 | 2.397 | 0.017 |
| Number of targets | 0.109 | 0.029 | 3.710 | 0.000 |
| Total path length | -0.078 | 0.021 | -3.737 | 0.000 |

| ***Final*** **Random effects engagement** | **variance** | **std. dev** | **corr** |
|---|---|---|---|
| Pair intercept | 0.05414 | 0.2327 | |
| Autoregressive covariance | 0.02112 | 0.1453 | -0.87 |
| Residual | 0.02660 | 0.1631 | |
| Number of observations: 275 | Number of pairs: 23 | | |

*Agreement*

| ***Initial*** **Predictor of agreement** | **estimate** | **std. error** | **t value** | **p value** |
|---|---|---|---|---|
| **Intercept** | **0.722** | **0.024** | **30.602** | **0.000** |
| Autoregressive covariance | -0.113 | 0.070 | -1.613 | 0.133 |
| Block (time) | -0.001 | 0.002 | -0.503 | 0.615 |
| Condition (same vs diff) | 0.005 | 0.008 | 0.619 | 0.537 |
| Neuroticism differences | 0.024 | 0.021 | 1.139 | 0.266 |
| **Extraversion differences** | **-0.045** | **0.021** | **-2.173** | **0.041** |
| Openness differences | 0.009 | 0.020 | 0.452 | 0.655 |
| Agreeableness differences | 0.017 | 0.023 | 0.748 | 0.462 |
| Conscientiousness differences | -0.019 | 0.020 | -0.937 | 0.358 |
| Autism quotient differences | 0.036 | 0.021 | 1.681 | 0.106 |
| Empathy differences | 0.016 | 0.020 | 0.805 | 0.429 |
| Distance covered by finger moves | 0.004 | 0.019 | 0.196 | 0.845 |
| Number of finger moves | 0.005 | 0.014 | 0.321 | 0.749 |
| Synchrony (WLCC-coef-mean) | -0.004 | 0.014 | -0.310 | 0.757 |
| Strength of relation (ppWLCC-coef-mean) | 0.005 | 0.018 | 0.295 | 0.769 |
| **Variability of relation (ppWLCC-coef-std)** | **0.015** | **0.008** | **1.885** | **0.061** |
| Time-lag (ppWLCC-lag-mean) | -0.011 | 0.013 | -0.802 | 0.424 |
| Switching (ppWLCC-lag-std) | 0.006 | 0.009 | 0.697 | 0.486 |
| Mutual information | 0.011 | 0.009 | 1.219 | 0.224 |
| Phase-slope index | -0.001 | 0.008 | -0.103 | 0.918 |
| **Number of targets** | **0.045** | **0.020** | **2.193** | **0.029** |
| Target sequence complexity | -0.014 | 0.013 | -1.026 | 0.306 |
| Time on obstacles | 0.006 | 0.020 | 0.316 | 0.752 |
| **Total path length** | **0.036** | **0.022** | **1.621** | **0.106** |

| ***Final*** **Predictor of agreement** | **estimate** | **std. error** | **t value** | **p value** |
|---|---|---|---|---|
| Intercept | 0.715 | 0.024 | 30.124 | 0.000 |
| Autoregressive covariance | -0.129 | 0.073 | -1.777 | 0.101 |
| Extraversion differences | -0.050 | 0.023 | -2.194 | 0.041 |
| Variability of relation | 0.015 | 0.007 | 1.971 | 0.050 |
| Number of targets | 0.059 | 0.016 | 3.704 | 0.000 |
| Total path length | 0.033 | 0.013 | 2.532 | 0.012 |

| ***Final*** **Random effects agreement** | **variance** | **std. dev** | **corr** |
|---|---|---|---|
| Pair intercept | 0.01208 | 0.1099 | |
| Autoregressive covariance | 0.04320 | 0.2078 | 0.46 |
| Residual | 0.01043 | 0.1021 | |
| Number of observations: 275 | Number of pairs: 23 | | |

*Predictability*

| ***Initial*** **Predictor of predictability** | **estimate** | **std. error** | **t value** | **p value** |
|---|---|---|---|---|
| **Intercept** | **0.738** | **0.026** | **28.611** | **0.000** |
| Autoregressive covariance | -0.046 | 0.074 | -0.624 | 0.543 |
| Block (time) | 0.000 | 0.002 | -0.245 | 0.807 |
| Condition (same vs diff) | 0.003 | 0.007 | 0.465 | 0.643 |
| Neuroticism differences | 0.014 | 0.019 | 0.761 | 0.455 |
| Extraversion differences | -0.028 | 0.019 | -1.453 | 0.160 |
| Openness differences | 0.026 | 0.019 | 1.413 | 0.171 |
| **Agreeableness differences** | **0.037** | **0.021** | **1.793** | **0.085** |
| **Conscientiousness differences** | **-0.036** | **0.018** | **-1.952** | **0.063** |
| Autism quotient differences | 0.029 | 0.020 | 1.477 | 0.153 |
| Empathy differences | 0.001 | 0.018 | 0.064 | 0.950 |
| Distance covered by finger moves | 0.005 | 0.018 | 0.267 | 0.789 |
| Number of finger moves | -0.001 | 0.013 | -0.069 | 0.945 |
| Synchrony (WLCC-coef-mean) | 0.002 | 0.013 | 0.132 | 0.895 |
| Strength of relation (ppWLCC-coef-mean) | 0.005 | 0.016 | 0.286 | 0.775 |
| **Variability of relation (ppWLCC-coef-std)** | **0.017** | **0.007** | **2.420** | **0.016** |
| Time-lag (ppWLCC-lag-mean) | -0.009 | 0.012 | -0.767 | 0.444 |
| Switching (ppWLCC-lag-std) | 0.000 | 0.008 | -0.020 | 0.984 |
| Mutual information | 0.002 | 0.003 | 0.682 | 0.496 |
| Phase-slope index | -0.226 | 0.302 | -0.747 | 0.456 |
| **Number of targets** | **0.049** | **0.018** | **2.713** | **0.007** |
| Target sequence complexity | -0.017 | 0.012 | -1.401 | 0.162 |
| Time on obstacles | -0.020 | 0.018 | -1.092 | 0.276 |
| Total path length | 0.000 | 0.020 | 0.018 | 0.986 |

| ***Final*** **Predictor of predictability** | **estimate** | **std. error** | **t value** | **p value** |
|---|---|---|---|---|
| Intercept | 0.741 | 0.019 | 38.637 | 0.000 |
| Autoregressive covariance | -0.038 | 0.076 | -0.493 | 0.630 |
| Agreeable differences | 0.047 | 0.019 | 2.424 | 0.024 |
| Conscientious differences | -0.040 | 0.019 | -2.087 | 0.049 |
| Variability of relation | 0.016 | 0.007 | 2.363 | 0.019 |
| Number of targets | 0.066 | 0.011 | 5.784 | 0.000 |
| Time on obstacles | -0.021 | 0.010 | -2.088 | 0.038 |

| ***Final*** **Random effects predictability** | **variance** | **std. dev** | **corr** |
|---|---|---|---|
| Pair intercept | 0.007763 | 0.08811 | |
| Autoregressive covariance | 0.054269 | 0.23296 | 0.21 |
| Residual | 0.008413 | 0.09172 | |
| Number of observations: 275 | Number of pairs: 23 | | |

### 2. Leave-one-out cross-validation of the final models' fixed effects

The cross-validation of fixed effects yielded highly significant and strong correlation between predicted and observed experience ratings, speaking to the generalisability of our findings - in the model of *engagement*, this is true for repeated measures (REP) (r = .454, p < .001), but not when aggregating over time (AVRG) (r = .12, p = .58), which is in line with the significant impact of the factor time on engagement. For the models of agreement and predictability, our cross validation yielded highly significant correlations between predicted and observed ratings, both for repeated measures as well as the pair average (*agreement*: (REP) r = .433, p < .001; (AVRG) r = .486, p = .019; *predictability:* (REP) r = .454, p < .001; (AVRG): r = .632, p = .001).

Note that the random intercept of the final models captured a greater part of the total variability in the model of engagement (67%), compared to agreement (54%) and predictability ratings (52%). Overall, generalisability hence appears higher for the latter two models.

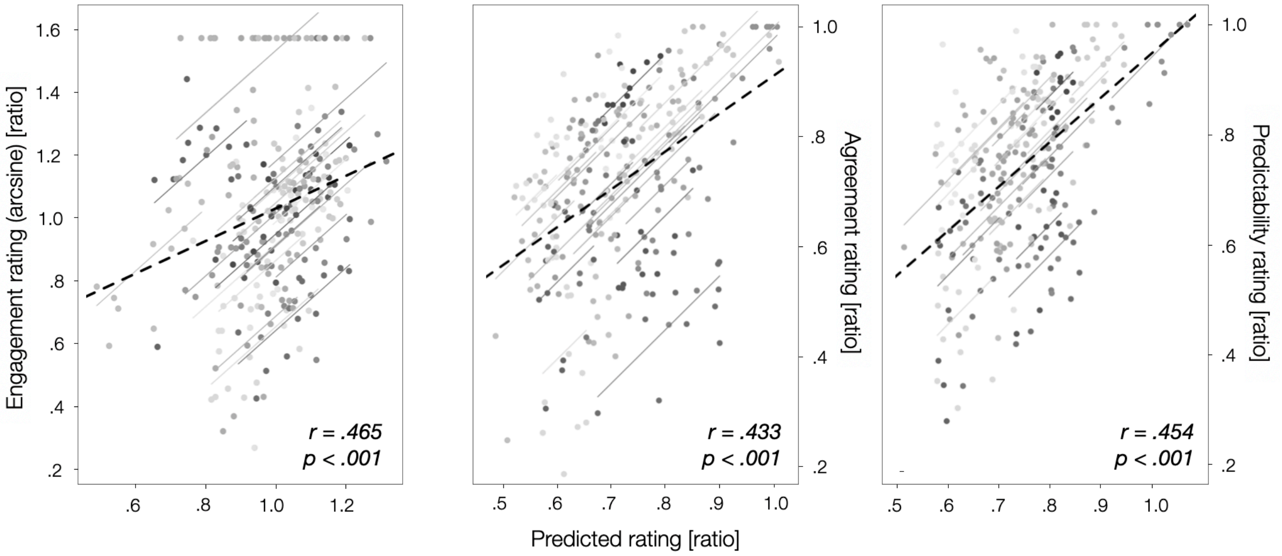


**Figure SC.1** Results from the leave-one-out cross-validation of the fixed effects in the final models of participants' engagement, agreement and predictability ratings (*left to right*). The plots show repeated measures correlations of observed (real) ratings with those predicted based on data from all but the present pair (12 x 2 values per pair). The dashed bar and statistics in the bottom right of the plot indicate the group average repeated measures fit - solid lines present individual pair fits.

### 3. Potential link between agreeableness differences and leader-follower roles (interpersonal time lag)

**Figure SC.2** Plotting agreeableness differences between players against their interpersonal time lag. The plot shows mean values of interpersonal time lag across all joint play trials, as well as the difference in the NEO-FFI agreeableness score between players. The line presents a second order polynomial fit through the data.

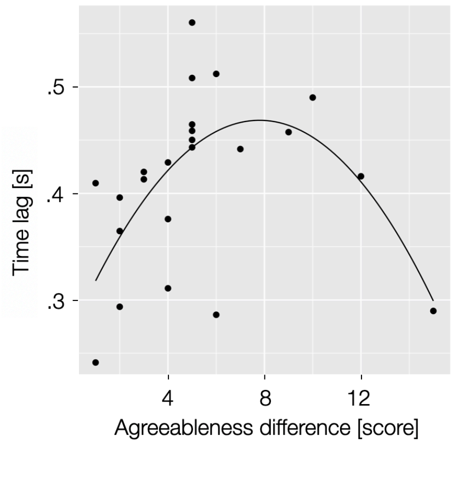

**Supplementary Materials D**

**Statistics for MANOVAs, individual measure ANOVAs, as well as post-hoc pairwise comparisons between individual blocks and trial segments**

**Tables SD.1** MANOVAs and Post hoc ANOVAs - main and two-way interaction effects. All p-values are FDR corrected at the MANOVA level, as well as within each family at the ANOVA level (here shown in separate tables). Significance levels are additionally indicated by *** (p < .001), ** (p < .01), * (p < .05), + (p < .10). The three-way interaction effect is not included in this table: all p > .18 and all MATS/ATS < 1.8.

| *MANOVAs* | Session | Condition | Block | Session by condition | Session by block | Condition by block |
|---|---|---|---|---|---|---|
| **Experience** | + p = .076, MATS = 13.93 | p = .153, MATS = 4.92 | **** p = .009, MATS = 10.09** | p = .754, MATS = .54 | p = .244, MATS = 4.01 | p = .238, MATS = 3.76 |
| **Gaming behaviour** | **** p = .009, MATS = 33.02** | + p = .087, MATS = 7.91 | ***** p < .001, MATS = 17.01** | p = .570, MATS = 1.25 | p = .378, MATS = 3.08 | p = .338, MATS = 3.74 |
| **Basic movement** | p = .509, MATS = .88 | p = .971, MATS = .02 | + p = .087, MATS = 3.51 | p = .461, MATS = .63 | p = .461, MATS = 1 | p = .813, MATS = .20 |
| **Movement coordination** | **** p = .006, MATS = 35.82** | ***** p < .001, MATS = 97.5** | p = .101, MATS = 20.06 | p = .143, MATS = 7.25 | p = .334, MATS = 12.8 | p = .207, MATS = 17.8 |

| *post hoc ANOVAs Exp* | Session | Condition | Block | Session by condition | Session by block | Condition by block |
|---|---|---|---|---|---|---|
| **Engagement** | + p = .054, ATS = 8.39 | p = .521, ATS = .63 | *** p = .017, ATS = 8.68** | p = .529, ATS = .54 | p = .975, ATS = .03 | p = .443, ATS = 1.25 |
| **Agreement** | p = .279, ATS = 2.15 | + p = .053, ATS = 6.51 | + p = .053, ATS = 5.76 | p = .523, ATS = .64 | p = .542, ATS = .72 | p = .446, ATS = 1.08 |
| **Predictability** | p = .159, ATS = 3.59 | p = .229, ATS = 2.44 | p = .156, ATS = 2.74 | p = .672, ATS = .27 | + p = .053, ATS = 4.92 | + p = .084, ATS = 3.98 |

| *post hoc ANOVAs Gam* | Session | Condition | Block | Session by condition | Session by block | Condition by block |
|---|---|---|---|---|---|---|
| **Targets** | ***** p < .001, ATS = 29.15** | *** p = .030, ATS = 6.81** | **** p = .003, ATS = 8.43** | p = .877, ATS = .03 | p = .478, ATS = 1.03 | p = .569, ATS = .67 |
| **Obstacle time** | *** p = .024, ATS = 7.17** | *** p = .046, ATS = 5.90** | **** p = .005, ATS = 9.08** | p = .569, ATS = .55 | p = .406, ATS = 1.25 | p = .242, ATS = 2.03 |
| **Path length** | **** p = .008, ATS = 10.30** | p = .559, ATS = .64 | ***** p < .001, ATS = 13.91** | p = .329, ATS = 1.46 | p = .451, ATS = 1.20 | p = .654, ATS = .49 |
| **Complexity** | p = .180, ATS = 2.77 | p = .520, ATS = .75 | p = .172, ATS = 2.32 | p = .983, ATS = 0 | p = .242, ATS = 2.03 | + p = .079, ATS = 3.75 |

| *post hoc ANOVAs Mov* | Session | Condition | Block | Session by condition | Session by block | Condition by block |
|---|---|---|---|---|---|---|
| **Number of moves** | p = .601, ATS = .61 | p = .914, ATS = .01 | + p = .068, ATS = 5.73 | p = .543, ATS = .75 | p = .788, ATS = .37 | p = .788, ATS = .34 |
| **Distance** | p = .383, ATS = 1.47 | p = .869, ATS = .06 | p = .383, ATS = 1.39 | p = .327, ATS = 2.06 | p = .229, ATS = 2.31 | p = .869, ATS = .16 |

| *post hoc ANOVAs MovC* | Session | Condition | Block | Session by condition | Session by block | Condition by block |
|---|---|---|---|---|---|---|
| **Synchrony** | ***** p < .001, ATS = 18.65** | ***** p < .001, ATS = 34.14** | p = .423, ATS = 1.43 | + p = .053, ATS = 7.39 | p = .515, ATS = 1.03 | p = .920, ATS = .26 |
| **Strength of relation** | ***** p < .001, ATS = 20.25** | ***** p < .001, ATS = 19.91** | p = .498, ATS = 1.22 | p = .292, ATS = 2.79 | p = .423, ATS = 1.37 | p = .971, ATS = .11 |
| **Stability of relation** | p = .125, ATS = 4.87 | p = .423, ATS = 1.44 | p = .579, ATS = .92 | p = .497, ATS = 1.05 | p = .184, ATS = 2.97 | p = .423, ATS = 1.45 |
| **Time lag** | + p = .084, ATS = 5.35 | p = .184, ATS = 3.56 | p = .372, ATS = 1.69 | p = .884, ATS = .15 | p = .995, ATS = .07 | p = .423, ATS = 1.63 |
| **Switching** | p = .278, ATS = 2.55 | p = .883, ATS = .14 | p = .928, ATS = .20 | p = .527, ATS = 1.02 | p = .813, ATS = .42 | p = .928, ATS = .20 |
| **Mutual Information** | p = .141, ATS = 4.14 | ***** p < .001, ATS = 40.55** | ***** p < .001, ATS = 13.03** | *** p = .046, ATS = 7.77** | p = .310, ATS = 2 | *** p = .020, ATS = 9.91** |
| **Phase slope index** | p = .141, ATS = 4.36 | p = .196, ATS = 3.5 | p = .813, ATS = .45 | p = .920, ATS = .07 | p = .920, ATS = .28 | p = .423, ATS = 1.47 |

**Table SD.2** Post hoc paired comparisons for significant and trending two-way interaction effects with block. p-values are FDR corrected within each family (together with the ANOVAs for the effects of session, condition and block, as well as post-hoc paired comparisons for individual blocks and trial segments). Significance levels are additionally indicated by *** (p < .001), ** (p < .01), * (p < .05), + (p < .10). The three-way interaction effect is not included in this table: all p > .18 and all MATS/ATS < 1.8.

| *post hoc paired comparisons ANOVA (interaction effects)* | Session by block (1-2) | Session by block (2-3) | Session by block (1-3) | Condition by block (1-2) | Condition by block (2-3) | Condition by block (1-3) |
|---|---|---|---|---|---|---|
| **Predictability** | *** p = .024, ATS = 10.93** | + p = .100, ATS = 4.89 | p = .545, ATS = .62 | + p = .072, ATS = 6.48 | p = .819, ATS = .12 | + p = .056, ATS = 7.65 |
| **Complexity** | x | x | x | p = .615, ATS = .39 | *** p = .044, ATS = 6.17** | *** p = .047, ATS = 5.30** |
| **Mutual Information** | x | x | x | p = .288, ATS = 2.65 | *** p = .021, ATS = 10.8** | *** p = .037, ATS = 9.83** |

x — Interaction effect not significant / no trend.

**Table SD.3** Post hoc ANOVAs - paired comparisons of individual blocks and trial segments for measures that show significant main effects of block or trial segment, respectively. p-values are FDR corrected within each family of observation - together with the ANOVAs for the effects of session, condition and block, as well as post-hoc paired comparisons for significant interaction effects. Significance levels are additionally indicated by *** (p < .001), ** (p < .01), * (p < .05), + (p < .10).

| *Post hoc paired comparison* ***ANOVAs*** | **Block 1 vs 2** | **Block 2 vs 3** | **Block 1 vs 3** | **Trial segm. 1 vs 2** | **Trial segm. 2 vs 3** | **Trial segm. 1 vs 3** |
|---|---|---|---|---|---|---|
| **Engagement** | p = .224, ATS = 2.64 | + p = .053, ATS = 6.37 | *** p = .017, ATS = 16.54** | x | x | x |
| **Agreement** | p = .975, ATS = 0 | *** p = .045, ATS = 8.37** | + p = .053, ATS = 6.67 | x | x | x |
| **Predictability** | p = .306, ATS = 1.77 | p = .458, ATS = .94 | + p = .084, ATS = 5.71 | x | x | x |
| **Targets** | p = .478, ATS = .91 | *** p = .030, ATS = 7.87** | ***** p < .001, ATS = 15.75** | ***** p < .001, ATS = 33.66** | ***** p < .001, ATS = 44.52** | *** p = .012, ATS = 8.83** |
| **Obstacle time** | p = .776, ATS = .14 | **** p = .005, ATS = 11.17** | ***** p < .001, ATS = 23.39** | + p = .098, ATS = 3.81 | ***** p < .001, ATS = 160.4** | ***** p < .001, ATS = 712.9** |
| **Path length** | p = .570, ATS = .50 | ***** p < .001, ATS = 21.21** | ***** p < .001, ATS = 22.63** | ***** p < .001, ATS = 123.3** | **** p = .006, ATS = 10.75** | ***** p < .001, ATS = 111.1** |
| **Complexity** | p = .312, ATS = 1.68 | + p = .083, ATS = 4.77 | p = .571, ATS = .49 | x | x | x |
| **Number of moves** | p = .318, ATS = 2.03 | **** p = .005, ATS = 14.53** | *** p = .042, ATS = 6.98** | ***** p < .001, ATS = 30.36** | + p = .090, ATS = 5.81 | *** p = .042, ATS = 7.69** |
| **Distance** | x | x | x | ***** p < .001, ATS = 36.33** | p = .383, ATS = 1.44 | ***** p < .001, ATS = 20.69** |
| **Synchrony** | x | x | x | p = .230, ATS = 2.92 | **** p = .006, ATS = 10.96** | **** p = .006, ATS = 11.76** |
| **Strength of relation** | x | x | x | p = .125, ATS = 4.78 | ***** p < .001, ATS = 14.85** | ***** p < .001, ATS = 18.94** |
| **Time lag** | x | x | x | *** p = .039, ATS = 6.95** | p = .299, ATS = 2.25 | ***** p < .001, ATS = 13.37** |
| **Mutual Information** | p = .813, ATS = .24 | ***** p < .001, ATS = 16.25** | ***** p < .001, ATS = 15.27** | x | x | x |

| | |
|---|---|
| x | Data not available at the level of the trial segment. |
| x | Main effect (of block / trial segm.) not significant / no trend. |

| *MANOVAs and post hoc ANOVAs* | Trial segment |
|---|---|
| **Gaming behaviour** | ***** p < .001, MATS = 203.13** |
| **Basic movement** | ***** p < .001, MATS = 3.9** |
| **Movement coordination** | ***** p < .001, MATS = 16.95** |
| **Targets** | ***** p < .001, ATS = 19.73** |
| **Obstacle time** | ***** p < .001, ATS = 155** |
| **Path length** | ***** p < .001, ATS = 102.81** |
| **Number of moves** | **** p = .005, ATS = 14.08** |
| **Distance** | ***** p < .001, ATS = 21.79** |
| **Synchrony** | **** p = .006, ATS = 8.51** |
| **Strength of relation** | ***** p < .001, ATS = 13.26** |
| **Stability of relation** | p = .997, ATS = .04 |
| **Time lag** | **** p = .006, ATS = 8.48** |
| **Switching** | p = .141, ATS = 3.62 |

**Table SD.4** MANOVA and post hoc ANOVAs results for the effect of trial segment. p-values are FDR corrected at the MANOVA level (together with the MANOVAs of session, condition and block), as well as within each family at the ANOVA level (together with the ANOVAs of session, condition and block). Additionally, significance levels are indicated by *** ($p < .001$), ** ($p < .01$).

**Supplementary Materials E**

**Additional visualisations of ANOVA results**

### 1. Main effects of block, session and condition

Main effects of block

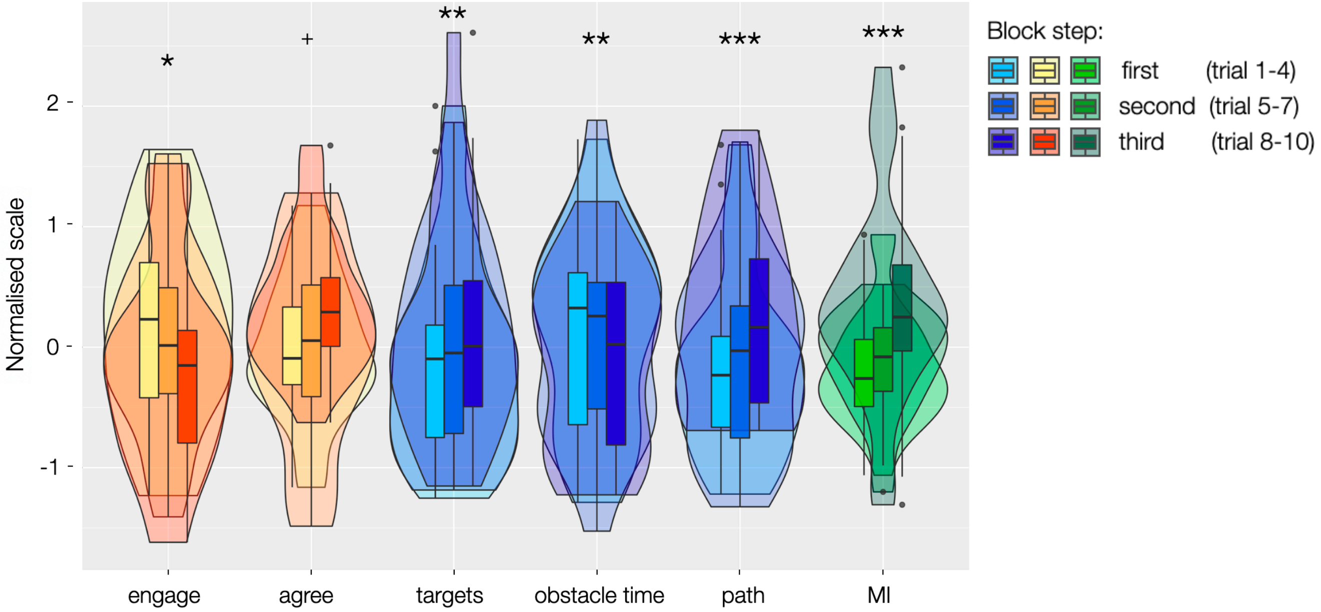


Main effects of session

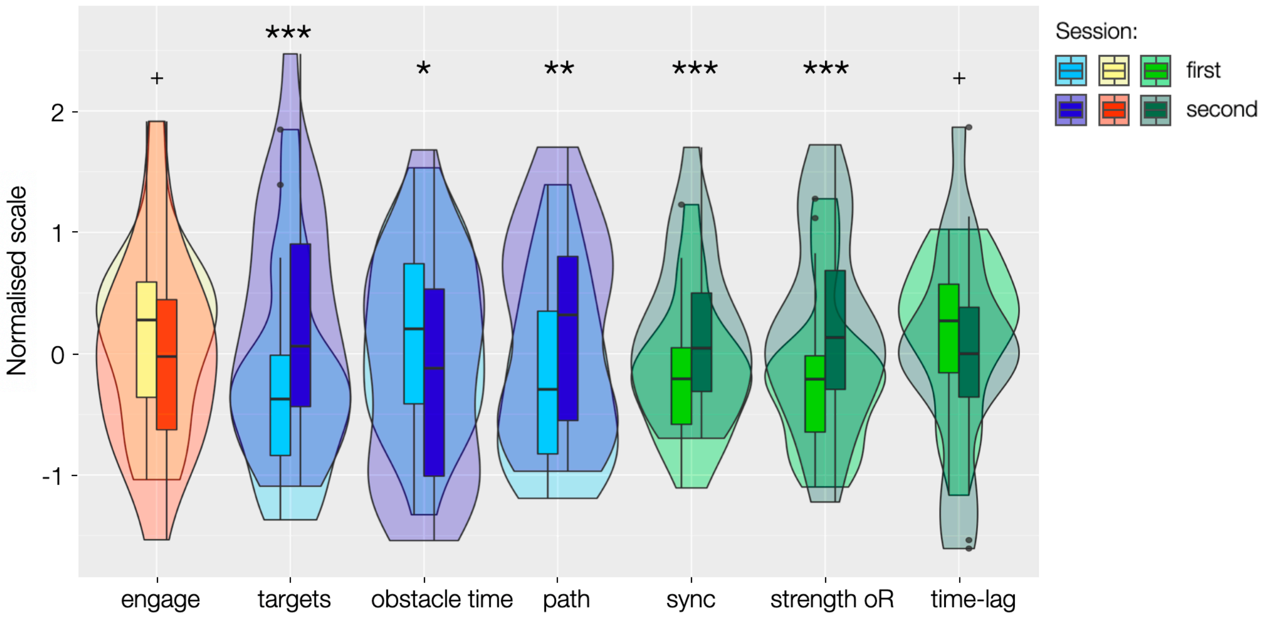


**Figure SE.1** Learning effects. Significant effects and trends over blocks (*top*) and sessions (*bottom*) All p-values are within-class of observation FDR corrected and indicated by *** ($p < .001$), ** ($p < .01$), * ($p < .05$), and + ($p < .10$).

Main effects of condition

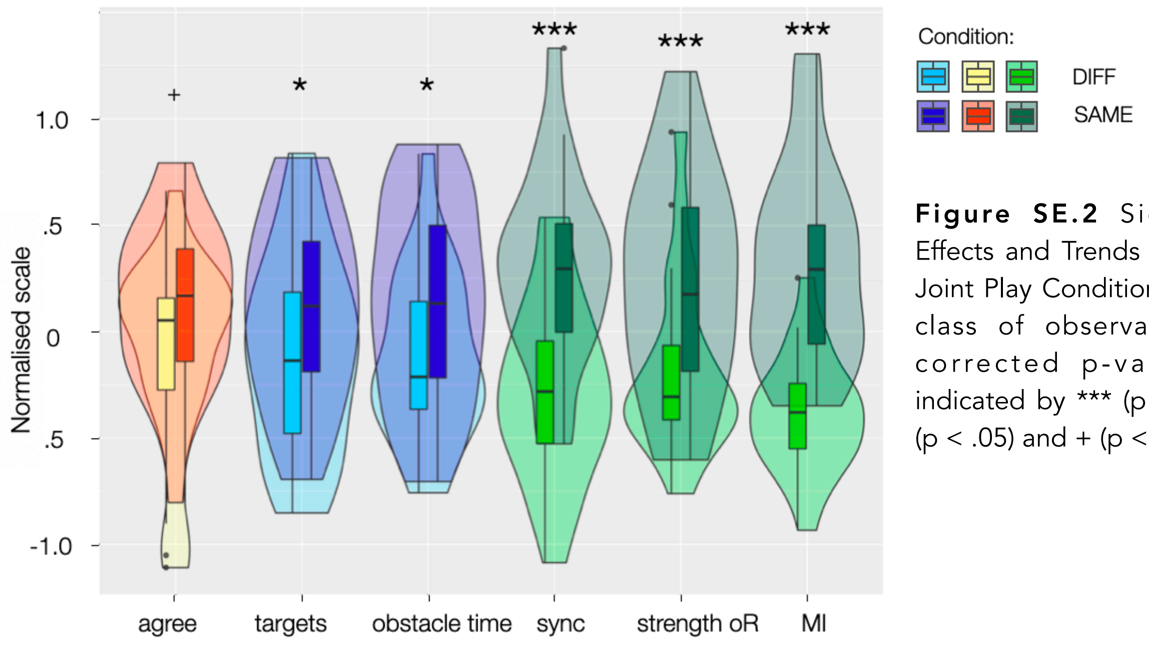


**Figure SE.2** Significant Effects and Trends across the Joint Play Conditions. Within-class of observation FDR corrected p-values are indicated by *** (p < .001), * (p < .05) and + (p < .10).

### 2. Main effects of trial segment (basic movement)

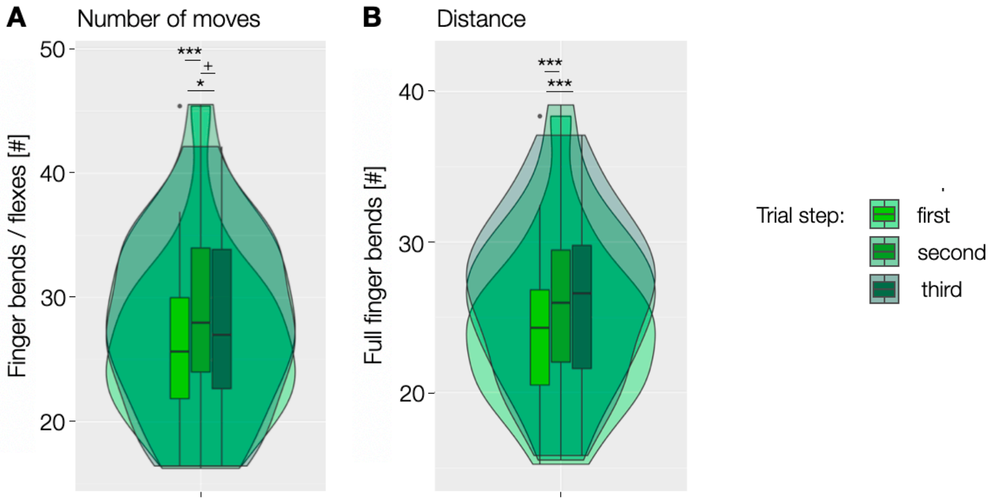


**Figure SE.3** Significant Effects of Trial Segment in Basic Movement and Movement Coordination Measures. Within-class of observation FDR corrected p-values are indicated by *** (p < .001), ** (p < .01), * (p < .05) and + (p < .10).

### 3. Additional interaction effects (trends)

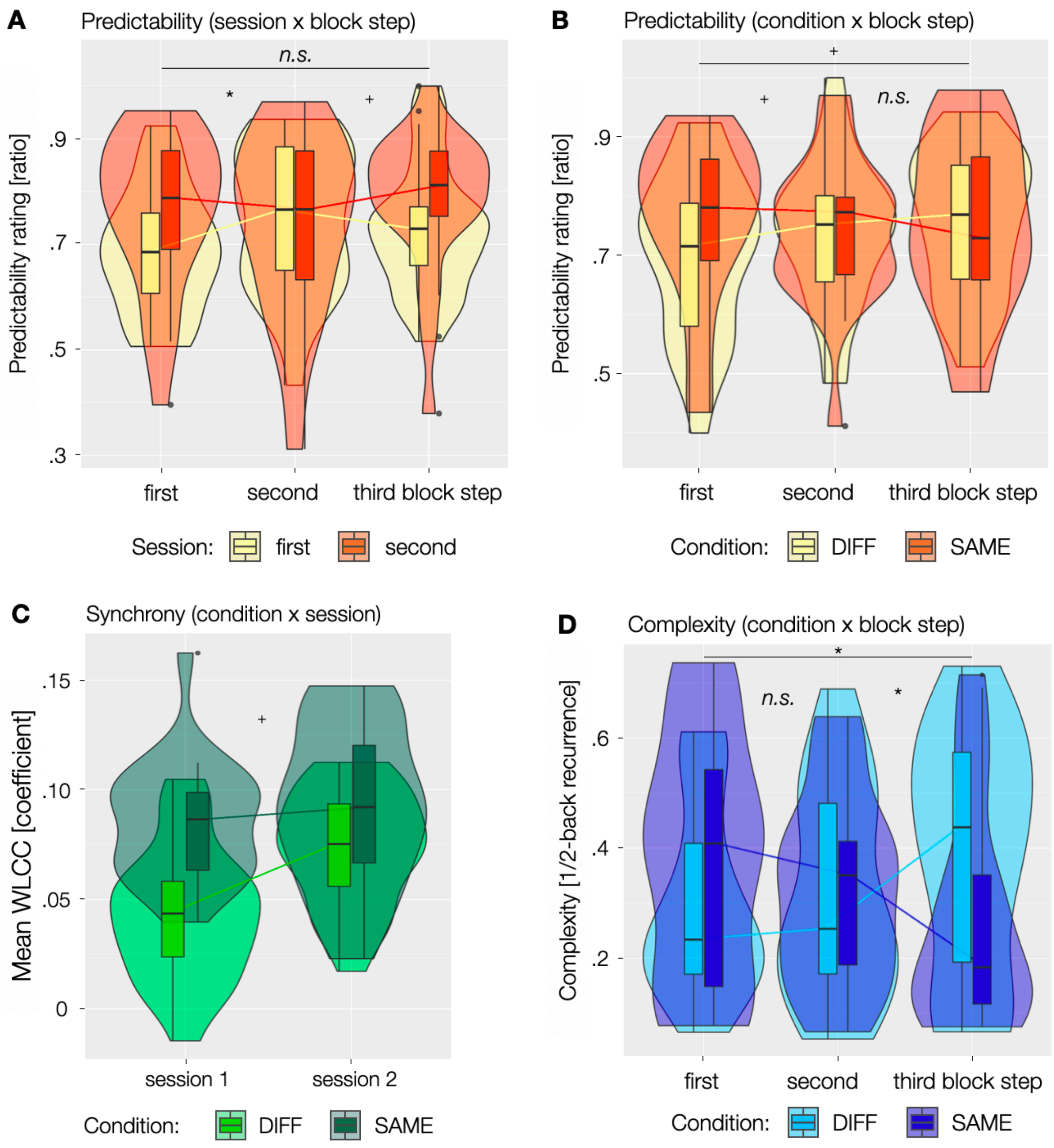


**Figure SE.4** Trends towards Interaction Effects. Within-family FDR corrected p-values are indicated by * ($p < .05$), + ($p < .10$) and *n.s.* ($p >= .10$).
